# Efficient and scalable modelling of cotranscriptional RNA folding with deterministic and iterative RNA structure sampling

**DOI:** 10.64898/2026.04.21.720041

**Authors:** Eliot Courtney, Edric K. Choi, Julius B. Lucks, Max Ward

**Affiliations:** Department of Computer Science & Software Engineering, The University of Western Australia, Western Australia, Australia; Interdisciplinary Biological Sciences Program, Northwestern University, Evanston, IL, USA; Center for Synthetic Biology, Northwestern University, Evanston, IL, USA; Department of Chemical and Biological Engineering, McCormick School of Engineering and Applied Science; Center for Water Research; Chemistry of Life Processes Institute; Northwestern University, Evanston, IL, USA

**Keywords:** RNA folding, RNA structure prediction, suboptimal folding, iterative sampling, cotranscriptional folding

## Abstract

RNA structure sampling is central to modelling RNA ensembles, yet stochastic sampling methods are non-exhaustive, scale poorly, and are biased towards low-free-energy structures, while current suboptimal folding approaches generate an unpredictable exponential number of structures. These limitations are particularly problematic for modelling cotranscriptional folding, where vectorial synthesis continuously reshapes the energy landscape during transcription, stabilising transient out-of-equilibrium structures. Here we introduce *iterative sampling*, a deterministic framework that enumerates unique RNA secondary structures in strict order of increasing free energy, enabling progressive and exhaustive exploration of the structure space up to an arbitrary stopping criterion. To implement this approach, we developed two scalable algorithms, iterative deepening and a persistent data structure approach, that incrementally traverse the expansion tree by evolving partial structures in place, avoiding redundant recomputation and fixed energy windows. Implemented in `memerna`, this approach achieves orders-of-magnitude speedups over existing tools (10x over `ViennaRNA`; 100x over `RNAstructure`). Integration within the sample-and-select framework (`R2D2`) improves structural diversity and identifies conformations with greater agreement with experimental data. Comprehensive sampling further enables direct comparison of equilibrium and cotranscriptionally restrained ensembles. Analysis of the resulting structural probability distributions uncovers kinetic traps and putative transcriptional pause sites, supporting an intuitive cotranscriptional folding mechanism in which local 3′-hairpin formation transiently stabilises upstream structure to delay large-scale rearrangement. Together, these results establish iterative sampling as a scalable and general framework for resolving out-of-equilibrium RNA cotranscriptional folding.

**Graphical abstract:** 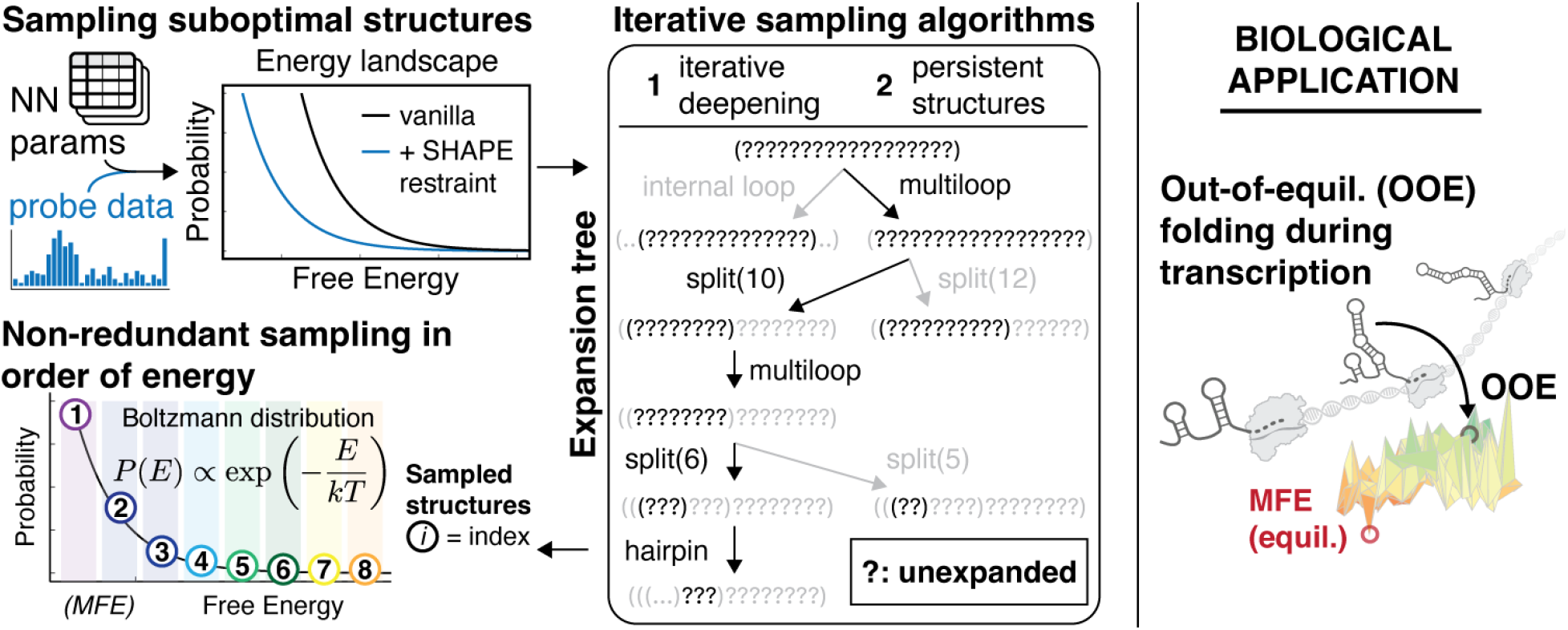

## 1 Introduction

Computational RNA structure sampling is a core component of nearly all RNA ensemble modelling approaches [1]. Ensemble modelling approaches have become a cornerstone of RNA biology and engineering, used for example to study multi-state RNA systems [2], uncover the impact of mutations on RNA structure [3], model out-of-equilibrium RNA folding pathways [4], and even to design new functional RNA [5]. All of these examples rely on RNA sampling algorithms to generate a set of representative candidate RNA structures out of an astronomically large number of possibilities [2]. These modelling methods are often based on stochastically sampling RNA structures based on their equilibrium Boltzmann probability, determined by calculating equilibrium partition functions [6]. While effective, this approach faces two limitations: (1) as the probability of sampling an RNA structure is equivalent to its Boltzmann weight, this approach is biased towards sampling structures with lower free energies; and (2) the stochastic framework is not prevented from sampling the same structure more than once, making it computationally inefficient to observe unique structures. As a result, existing approaches cannot explore the broader RNA folding landscape, with rare, high-energy, or kinetically relevant conformations systematically unrepresented or absent, despite their functional importance [7].

In addition to stochastic sampling, a variety of suboptimal folding approaches have been developed to broaden coverage beyond minimum-free-energy predictions [8]–[10], yet all remain restrained by how structure space is explored. Early work by Zuker and colleagues introduced heuristics to generate non-MFE structures by enforcing specific base-pair restraints and recomputing the optimal fold under those restrictions [9]. Although initially thought to provide broad sampling of suboptimal structures, these approaches do not systematically enumerate all alternatives and primarily probe local deviations from the MFE structure. Wuchty *et al*. [10] subsequently introduced a systematic algorithm that enumerates all structures within a predefined energy window of the MFE by exploring every path in the dynamic programming recursions and pruning some, based on the general dynamic programming suboptimal traceback finding algorithm of Waterman and Byers [11]. While formally complete within a specified energy window, this strategy becomes computationally intractable as the energy window expands, and neither of these approaches samples structures according to their Boltzmann probabilities. To address this limitation, Ding and Lawrence [6] developed stochastic sampling methods that draw structures from the Boltzmann ensemble, later refined for computational efficiency [12]. Although thermodynamically principled, stochastic sampling is inherently redundant and biased toward low-energy states, whereas classical suboptimal sampling is rigidly bounded by a fixed energy window. Notably, none of these approaches naturally support sampling structures in ranked order or continuing exploration until an arbitrary stopping criterion is met.

These limitations are particularly acute for studying cotranscriptional RNA folding, which often proceeds along nonequilibrium trajectories characterised by kinetic barriers and large-scale dynamic structural rearrangements [13]– [16]. During cotranscriptional folding, each additional nucleotide reshapes the energy landscape [7], allowing transient intermediates and kinetic traps to form that may never dominate at equilibrium. Experimental advances, including cotranscriptional structure probing, now provide nucleotide-resolution views of these dynamic processes [17]–[21]. Recently, a number of algorithms have been developed that can leverage this data to make experimentally restrained models of cotranscriptional folding. One such approach, named Reconstructing RNA Dynamics from Data (`R2D2`), employs a sample-and-select framework in which large sets of candidate secondary structures are first generated by stochastic Boltzmann sampling and then scored against experimental reactivity data to identify structures most consistent with the observations [4]. More broadly, related thermodynamics-dependent approaches combine data-directed sampling, clustering, or ensemble reweighting, using probing data to restrain or redistribute probabilities over thermodynamically generated structure ensembles [2]. In all of these cases, inadequate sampling does not merely reduce accuracy, it fundamentally limits the types of structures that are analysed in these approaches.

Here, we address these limitations and introduce a new sampling algorithm that enables iterative sampling, which we refer to as generating sequences in sorted order one at a time until an arbitrary stopping condition is reached. This approach maximises the number of unique samples subject to any predicate, such as computational budget, a structure diversity threshold, or the traditionally used energy delta. Enumerating RNA secondary structures in order of increasing free energy enables computationally efficient, progressive exploration of the RNA folding landscape. We present two complementary algorithms for suboptimal folding that support this strategy, an iterative deepening based approach [22] extending the algorithm of Wuchty *et al*., enabling many additional optimisations, and a novel approach based on persistent data structures [23], each offering distinct trade-offs between memory usage and computational efficiency. In particular, our iterative deepening based approach uses minimal memory but producing the next suboptimal structure may have higher latency. Our persistent data structure based approach uses more memory but has much less variance in the amount of time it takes to produce the next suboptimal structure. We also do a new analysis of the distribution of RNA structures versus their energy and shared substructures, showing that the distribution is locally exponential but globally normal, and that sampling structures in the exponential regime uses only a linearly growing set of shared substructures. Rather than treating sampling as a probabilistic approximation of equilibrium, our framework makes structural diversity, convergence, and rare states explicitly observable. These methods are implemented in the existing RNA folding package `memerna` [24], and together span a continuum of possible strategies between time- and space-efficient sampling. We first develop these approaches and show that they are faster than both `ViennaRNA` and `RNAstructure` while finding structures that `ViennaRNA` and `RNAstructure` do not. Next we integrate this approach within `R2D2` to study cotranscriptional folding using experimental structure probing data, asking whether deterministic, non-redundant enumeration can expand structural coverage, improve reproducibility, and enable direct comparison of equilibrium and cotranscriptional folding landscapes. We show that even basic integration of `memerna` iterative sampling improves performance and accuracy. We further investigate incorporating chemical probing reactivities as pseudo-free energy restraints within iterative sampling, and show that iterative sampling comprehensively samples, allowing comparison of the underlying structure probability distributions across conditions. Overall, this work establishes iterative sampling as both a scalable algorithmic advance and a biologically meaningful framework for studying cotranscriptional RNA folding.

In Section 2 we review and analyse the existing algorithms for suboptimal folding and introduce our two new algorithms. In Section 4 we benchmark the time and memory performance of our suboptimal folding algorithms against existing implementations in `RNAstructure` [25] and `ViennaRNA` [26]. We also show the application as a sampling engine for `R2D2` and demonstrate a significant improvement in sampling efficiency, runtime, and sample distribution fitness.

## 2 Algorithmic discussion

Development of computational approaches for generating suboptimal RNA secondary structures generally fall into three approaches, including heuristic suboptimal enumeration methods, exhaustive enumeration within a fixed energy window, and stochastic sampling from a Boltzmann ensemble.

The first approach derives directly from the original Zuker-Stiegler algorithm [8]. Zuker [9] introduced a heuristic method for identifying suboptimal structures by finding the MFE structure enclosed by and exterior to each possible base pair (*i, j*). This algorithm finds at most 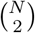 suboptimal structures which contain each possible base pair. While computationally efficient, this approach does not systematically sample all suboptimal structures. We describe Zuker’s suboptimal folding method in Supplementary Section C.1.1.

A second approach attempts to overcome this by exhaustively sampling within a fixed free energy window. The algorithm introduced by Wuchty *et al*. [10] performs a depth-first search of the directed acyclic graph formed by the dynamic programming recurrence relations, where states that exceed a specified energy delta are not traversed, enabling sampling of all suboptimal structures within a pre-defined energy window. However, selecting an appropriate energy window can be difficult in practice. This is particularly problematic when chemical probing pseudo-free energies make energy quantities difficult to interpret, or when the user wants to iterate through structures until a stopping condition is reached. The latter can be mitigated by iteratively increasing the energy window, but this comes with a performance cost. We introduce a variation of this algorithm that significantly improves performance of the iterative algorithm variant. We describe the algorithm of Wuchty *et al*. in Supplementary Section C.1.2, algorithm B, and Supplementary Table G.

The third approach is the stochastic sampling method of Ding and Lawrence [6]. The time complexity of this approach was later improved by Ponty [12], taking the worst case runtime from *O*(*N* ^2^) to *O*(*N* log *N*) and the average case runtime from 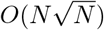 to *O*(*N* log *N*). The Ding and Lawrence algorithm stochastically samples structures according to their probability in the Boltzmann ensemble of structures as defined by McCaskill’s partition function algorithm [27]. This is useful when a statistical sample is needed, but is not suitable when non-redundant sampling is more suitable, as is often the case when searching for out-of-equilibrium folds. The LinearSampling algorithm by Zhang *et al*. [28] adapts this using beam search and caching to heuristically sample structures more quickly. We describe stochastic sampling in Supplementary Section C.1.3.

For the rest of this paper we will refer to suboptimal folding specifically as approaches similar to that of Wuchty *et al*. involving graph traversals of the dynamic programming recurrence relations and we will refer to stochastic sampling as approaches similar to that of Ding and Lawrence.

In Table 3 we compare algorithmic features of `memerna` with `RNAstructure` and `ViennaRNA`. In Table 2 we summarise `memerna` specific optimisations.

**Table 1:**
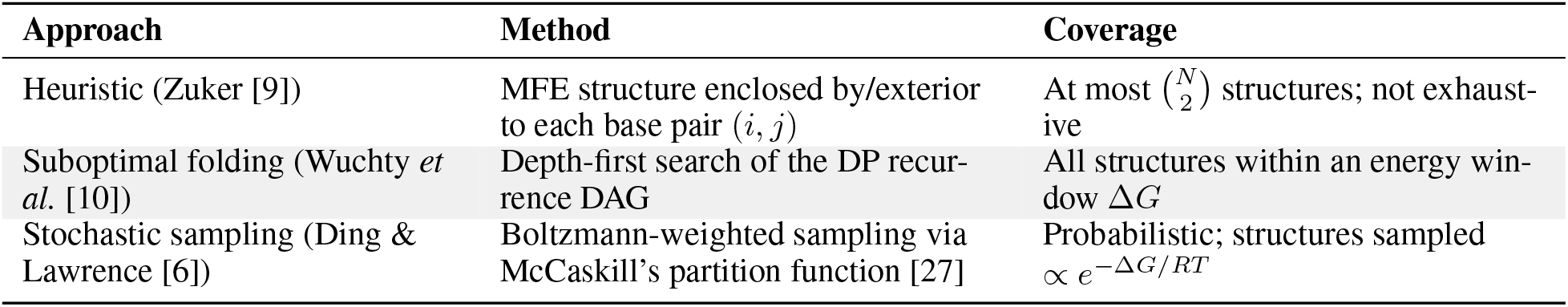
Summary of prior suboptimal RNA folding approaches.

**Table 2:** memerna-specific suboptimal folding optimisations.

| Optimisation | memerna iterative | memerna persistent |
| --- | --- | --- |
| Fast partial structures | Single structure updated in place during DFS | $O(1)$ memory PFS nodes |
| Expansion memoisation | Cached and sorted expansions per DP state with $O(1)$ lookup | Same |
| Next best expansion processing | Each visited state processes only its next best expansion | Same |
| Expansion pruning | Expansion processing pruned when exceeds energy delta | PFS naturally visits states in energy order |
| Sorted output generation | Iterative deepening on each discrete energy level, skipping empty energy levels | Inherent to PFS |
| Non-sorted fast path | Runs without iterative deepening for faster unsorted output | N/A |
| Depth first behaviour | Inherent to DFS | Monotonic node index in priority queue tie-breaks to deeper nodes |
| Memory cap on expansion cache | lowmem variant limits expansion-cache memory to $O(N^2)$ | Same |

**Table 3:** Algorithmic optimisation comparison. *S* denotes the number of output structures and *N* denotes the sequence length. Note that tree-state memory excludes the dynamic programming tables and expansion cache, which is *O*(*N* ^3^) for `memerna`, or *O*(*N* ^2^) for `memerna-lowmem` which is the LRU cache variant.

| Optimisation | ViennaRNA | RNAstructure | memerna iterative | memerna persistent |
| --- | --- | --- | --- | --- |
| Iterator capability | No | No | Yes (but may run the next DFS iteration) | Yes |
| Tree-state memory complexity | $O(SN)$ | $O(SN)$ | $O(N)$ | $O(S)$ |
| Memory per expansion | $O(N)$ copy | $O(N)$ copy | $O(1)$ incremental apply/undo | $O(1)$ via persistent data structure |
| Time per expansion | $O(N)$ copy | $O(N)$ copy | $O(1)$ incremental apply/undo | $O(1)$ new node with parent pointer |
| Process one expansion at a time | No | No | Yes | Yes |
| CTD handling | d2 natively; d1 and d3 by post-processing (misses structures) | Full | Full | Full |
| Algorithm kind | Wuchty *et al.* | Wuchty *et al.* | Wuchty *et al.*-like iterative deepening | Priority first search |

### 2.1 Our new approaches

We want to be able to repeatedly sample the next best suboptimal structure until an arbitrary stopping criterion is reached. One way to do this is to use a priority queue to perform a priority first search (PFS) of the *expansion tree* (see Section 2.2). There are a few issues with doing this naively:

- It takes a lot of memory to store all the partial structures on the priority queue.
- It is expensive in time and memory to enumerate all expansions at a node.

Another way to do this is to run the algorithm of Wuchty *et al*. multiple times (iterative deepening). It also has issues:

- It takes a lot of memory to store all the partial structures on the explicit stack.
- The output of the algorithm of Wuchty *et al*. is not ordered, and it is only trivial to sort it if it can fit in memory.
- It is hard to know what to set the energy delta to avoid producing too many structures.

In the next sections we describe how we enable both of these approaches to run performantly. Then in Section 4 we compare their performance in practice.

### 2.2 Definitions

Firstly, we have some definitions of concepts and functions used in the following explanation and pseudocode.

#### Suboptimal structure

A secondary structure with a free energy more than the minimum free energy (MFE). For brevity and consistency we also consider the MFE structure to be a suboptimal structure.

#### State

A state in the dynamic programming recurrence [6], [8], [10]. For example, (*st, en, a*) where *st* is the start index, *en* is the end index, and *a* indicates the dynamic programming table (for example, the *P*, *U*, *EXT*, or *U* 2 table [24] — the value here is dependent on the exact recursions used). A state covers a span of contiguous nucleotides, for example, *st* to *en*, inclusive.

#### Partial structure

A list of base pairs, CTD tags, a set of unexpanded states, and the current energy delta from the MFE. This represents a partially determined secondary structure. In the process of constructing suboptimal tracebacks, partial structures are gradually evolved towards complete structures. A complete structure will have no unexpanded states. In prose and pseudocode, we omit the explicit handling of CTD tags.

#### Expansion

A structure representing how to expand a partial structure at a particular state. For example, it may contain: Which child states are used by this state, which base pairs are added, which CTD tags are added, and the energy delta this expansion would add.

#### Expansion tree

The tree of partial structures. Edges in the tree correspond to expansions. Internal nodes correspond to partial structures. Leaf nodes correspond to suboptimal structures (that is, secondary structures).

#### MfeFold(*r*)

The Zuker-Stiegler algorithm [8] for computing the minimum free energy secondary structure. It runs in *O*(*N* ^3^) time and *O*(*N* ^2^) auxiliary memory. It returns the dynamic programming tables.

#### Partition(*r*)

The McCaskill algorithm [27] for computing the partition function. It runs in *O*(*N* ^3^) time and *O*(*N* ^2^) auxiliary memory. It returns the partition function dynamic programming tables.

#### Traceback(*mfe_dp_tables*)

Performs the optimal traceback given the set of dynamic programming tables. It runs in *O*(*N* ^2^) time and *O*(*N*) auxiliary memory. It returns a secondary structure.

#### Expand(*mfe_dp_tables, state*)

Generates all expansions for a given state. It runs in *O*(*N*) time and *O*(*N*) auxiliary memory. It returns a list of expansions.

*δ* - The difference from the minimum free energy (MFE) that all suboptimal structures should be less than or equal to.

*r* - The RNA primary structure.

#### 2.2.1 Expansion tree optimisations

In this section we describe improvements to expansion tree handling that apply to both the PFS approach and the iterative deepening approach.

It is only necessary to process one expansion when visiting a state, as long as it is the next best expansion for that state. The algorithm of Wuchty *et al*. by contrast iterates through every expansion while processing a state and adds it to the stack. In order to do this optimisation, we need to sort expansions by their energy delta compared to the best energy for that state. It is useful to avoid processing every expansion at each state because we are generally enumerating suboptimal structures in order from lowest to highest energy or only outputting structures within a certain delta from the MFE. So, we can avoid processing expansions that make the energy too high or only process the next best expansion. We describe this in Algorithm 1.

It is possible to map states to a linear integer index in constant time so lookup in *cache* can be done in constant time.

Storing *cache* can take *O*(*N* ^3^) memory because there are *O*(*N* ^2^) unique states and each state can have *O*(*N*) expansions (Corollary C.1.3). It is possible to reduce this to *O*(*N* ^2^) using a LRU cache — this is what `memerna-lowmem` referred to in the Supplementary Material does.

##### Algorithm 1

Pseudocode describing cached computation of sorted expansions. The cache can be made into an LRU cache to limit memory usage to *O*(*N* ^2^).

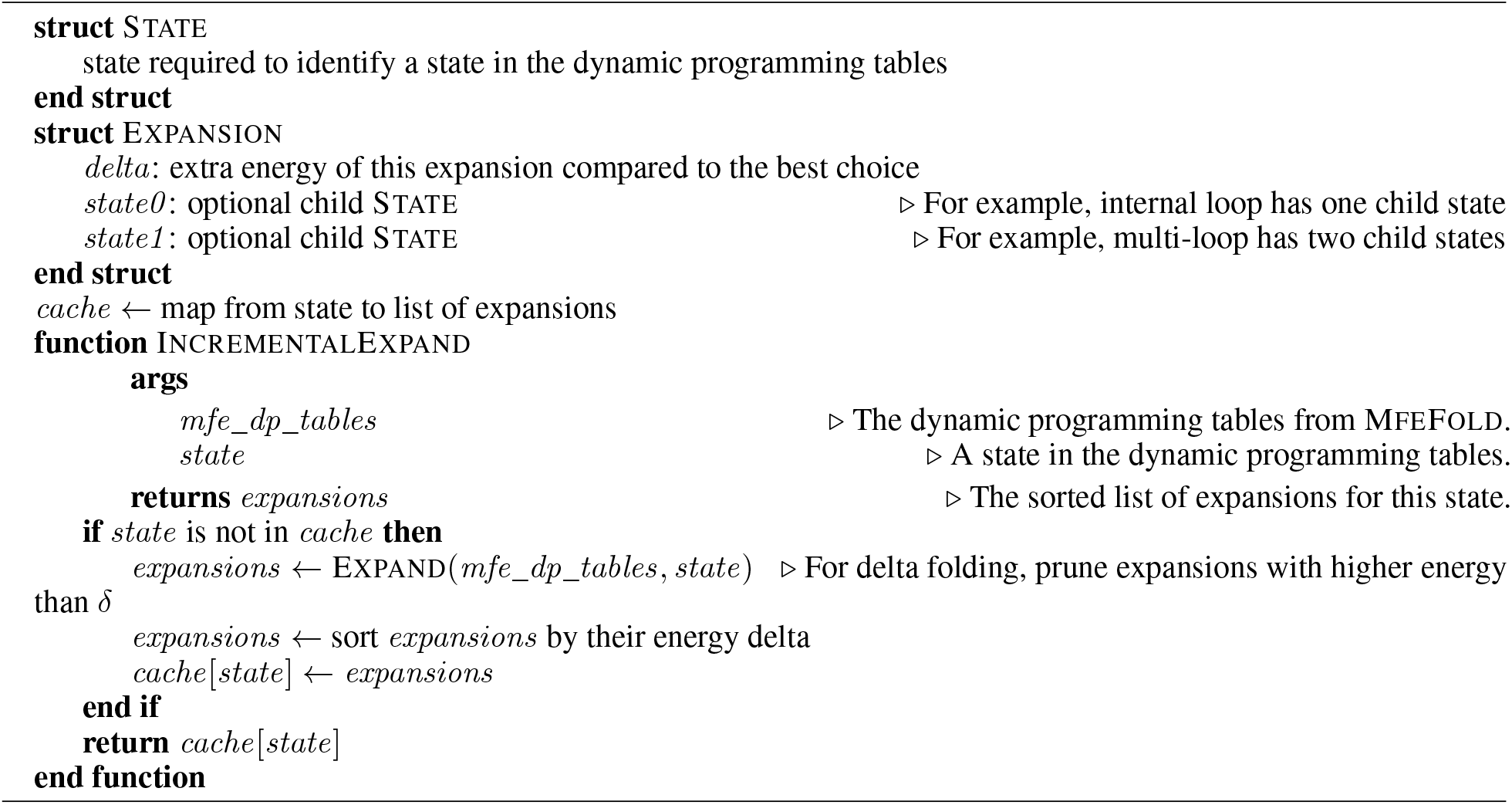

#### 2.2.2 Iterative suboptimal folding

We describe the iterative deepening version of the algorithm in Algorithm 2 and Algorithm E. Essentially, it is based on the algorithm of Wuchty *et al*. with extensive optimisations and modifications to allow iterative deepening to work well.

An issue with the algorithm of Wuchty *et al*. is that it stores entire partial structures on its stack and copies them for each new partial structure created. Since we are performing a DFS we can store the partial structure just once and update it incrementally as we traverse the expansion tree. This avoids the linear time copies for each expansion step. In combination with processing only a single expansion at a time using INCREMENTALEXPAND, we can visit a state in constant time. The pseudocode in Algorithm 2 shows the update of the base pairs of the partial structure (PartialStructure.*base_pairs*), but updating coaxial stacking, dangling ends, and terminal mismatch (CTDs) information can be handled in a similar way [24], [29].

To visit a state, we look at the next NODE on the top of the stack. We get the list of sorted expansions at the state *to_exp* using IncrementalExpand. We know what the next one to process by checking *exp_idx*.

There is a complication to deal with while processing an expansion. An expansion can have multiple child states that need to be visited (at most two in Turner 2004, since multi-loops are partitioned into a left and right sub-problem). One child state can be processed directly as the next node in the DFS. But the other needs to be saved (in the list *unexp* referred to in Algorithm 2) to be processed later, otherwise we would have child state remain unprocessed.

We handle this by popping unprocessed states from *unexp* when we run out of child states to process in the DFS. These states are special because we need to remember to re-insert them into *unexp* when going up to the parent and un-doing their effect on the incremental state. This is because we still have applied the expansion for this state somewhere higher up in the DFS tree, so until we backtrace up the DFS tree and remove that node, we must not forget to apply this state in any part of the DFS below that. We remember to do this by storing a boolean *should* _*unexp* in NODE for these special states.

In Figure 1 we show an example of how the DFS evolves a partial structure, and how the DFS state is updated incrementally as we traverse the expansion tree. In particular, we show how the unexpanded states are handled.

**Figure 1:**
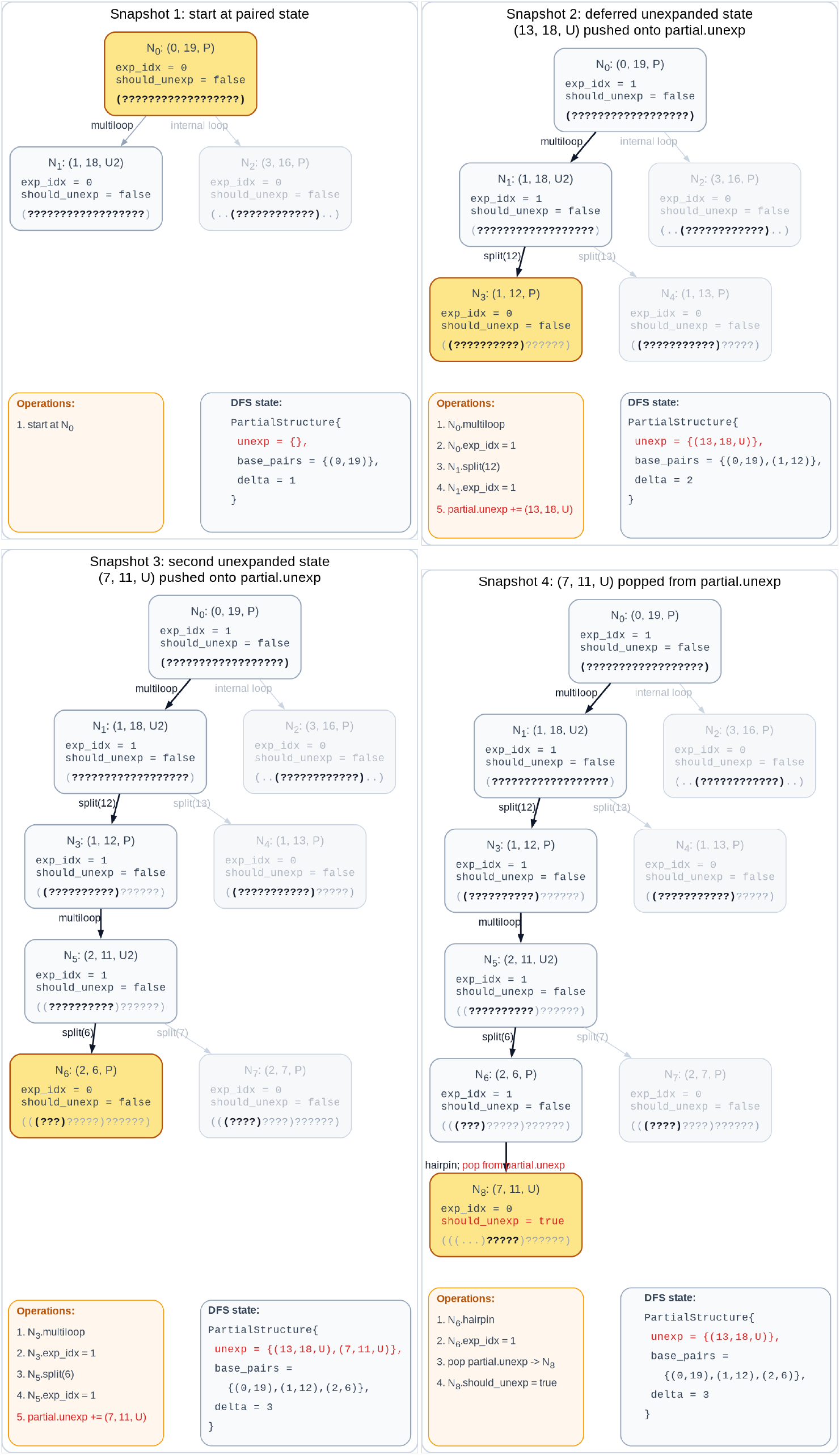
We illustrate the evolution of a partial structure in the iterative deepening suboptimal folding algorithm. Each snapshot shows what the DFS tree looks like at a particular point. Items shown in red are related to the handling of unexpanded states. Here we use the number of base pairs (Nussinov-like [30]) as the delta value, instead of difference to MFE, for simplicity.

By doing this, we get an *O*(1) per expansion implementation of suboptimal folding. By adding iterative deepening [22], we can support sorted output, outputting a specific number of structures rather than using an energy delta, or running for (approximately) a given time. Since the energy values are quantised, we perform the DFS to get everything with energy delta 0, then 0.1 (if using 0.1 precision), and so on. This is different to simply running an existing implementation of the algorithm of Wuchty *et al*. multiple times with increasing energy windows. Firstly, we keep track of the next energy delta level available to avoid re-running the DFS when it will not produce any new structures. Secondly, we only output structures at exactly each energy delta. Thirdly, we can stop producing structures at any point, even in the middle of an energy delta level (iterator functionality). Due to the exponential growth of the number of structures with energy it may not be feasible to fully enumerate the next energy delta level like would be required with existing approaches.

With this, we enable sorted output while avoiding regenerating structures that were already generated at previous energy levels. This also gives iterator-like support for iterative sampling because the DFS can be stopped whenever we no longer need more structures. The limitation is continuing to the next structure may require starting the next iterative-deepening pass at the next energy level.

##### Algorithm 2

Pseudocode describing a single step in iterative deepening suboptimal folding.

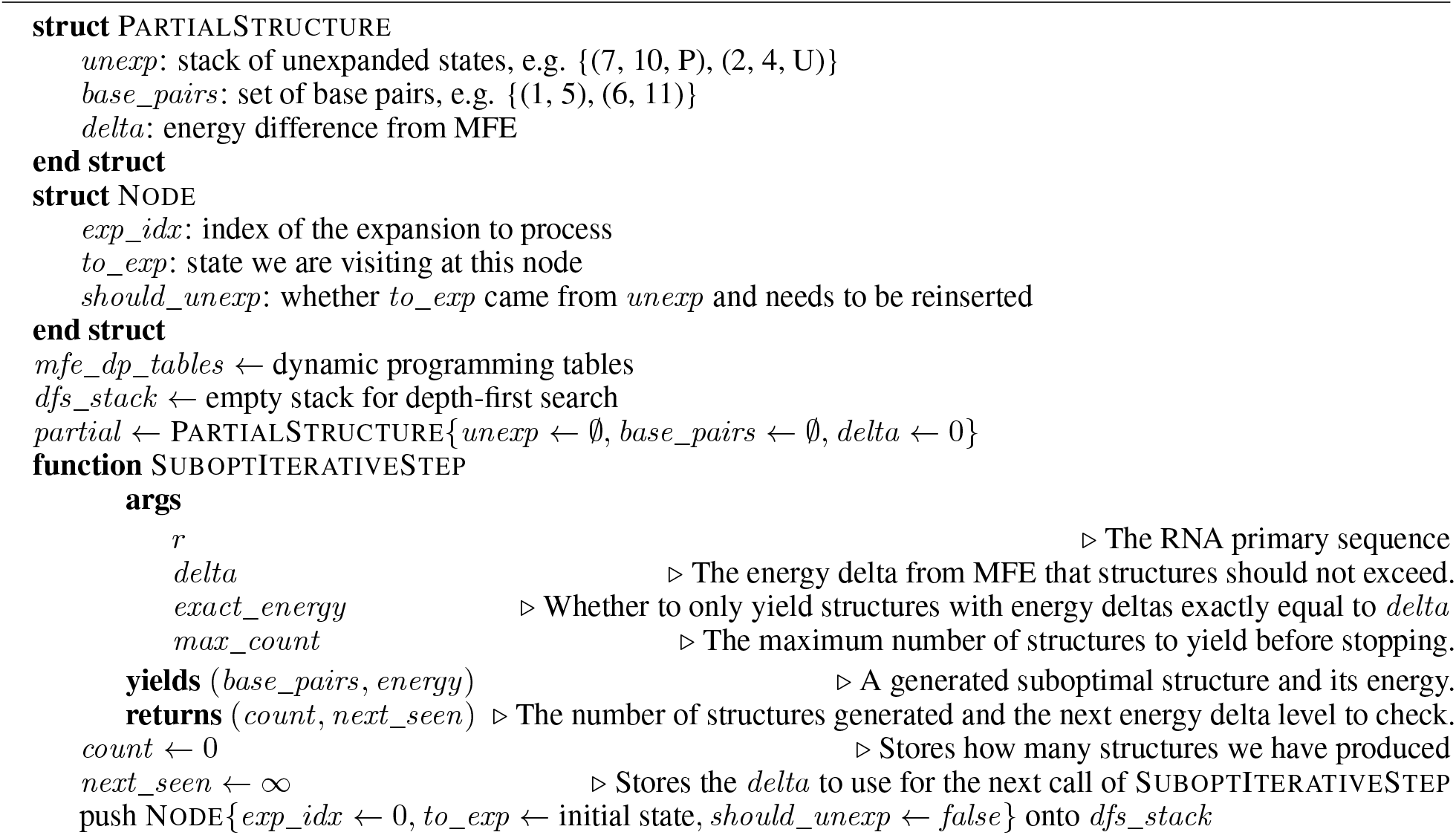
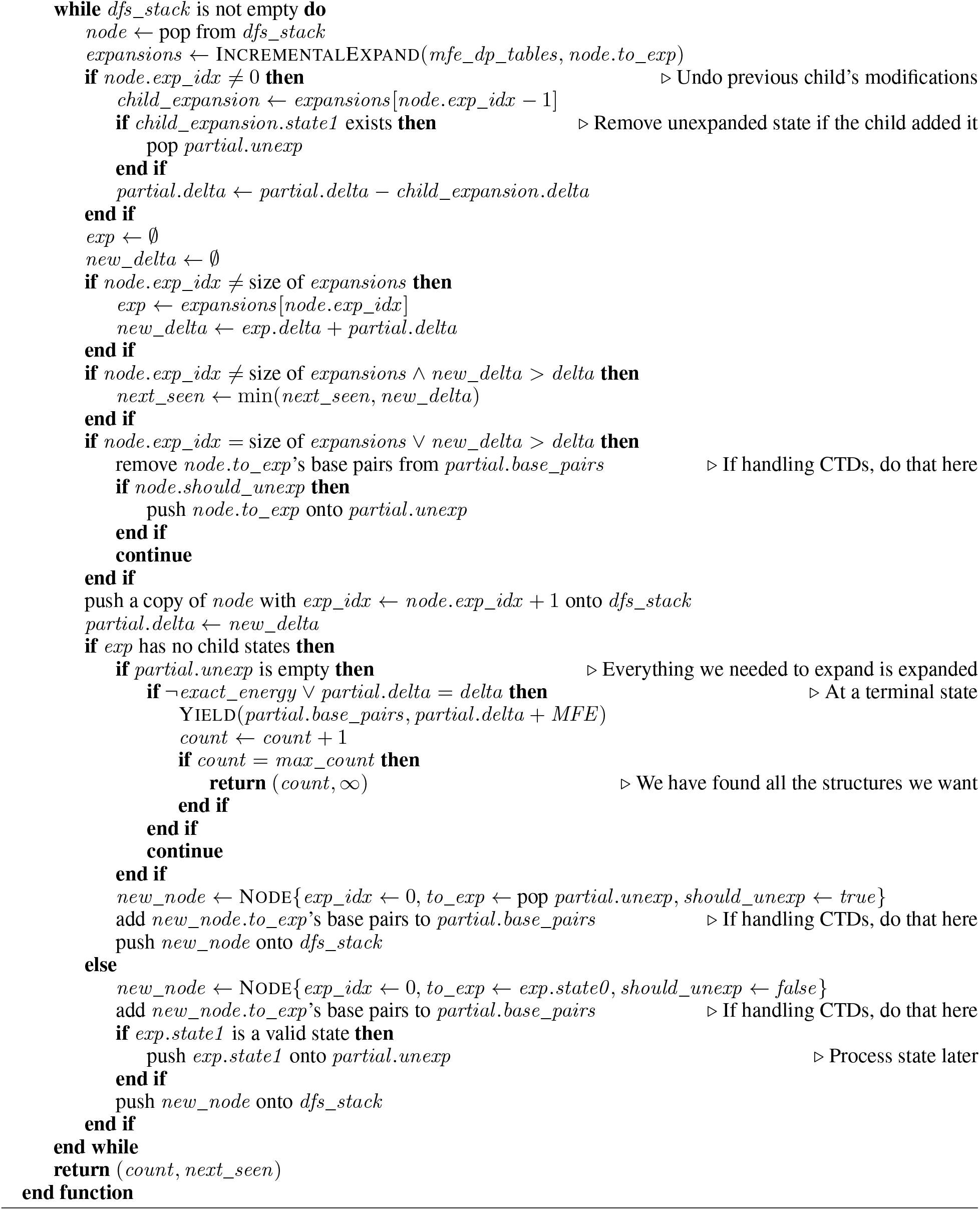

#### 2.2.3 Persistent suboptimal folding

The second algorithm we have is an improvement on the PFS approach by using a persistent data structure to save memory. In the iterative deepening approach we saved memory by evolving and unevolving the partial structure in place. Instead here we observe that the partial structures can be stored implicitly as a series of expansions to the empty partial structure. We keep a tree of expansions and construct each suboptimal structure by walking the tree from leaf node (representing a complete partial structure) to root node. See Algorithm D in the Supplementary Material for how this is done. The advantage of this approach over iterative deepening is that the time it takes to produce the next structure is small. Iterative deepening may need to start a new DFS which may take some time before it produces the next structure. This approach also does not rely on exponential growth of the number of structures with the energy delta level to be efficient. The trade-off is that this approach needs to store an *O*(*S*) amount of memory.

Simply creating a node for each expansion at a state and pushing it onto the priority queue would work but wastes memory in the same way that the algorithm of Wuchty *et al*. does by processing each expansion and pushing them all onto the stack. Instead, to process each expansion in order from lowest to highest energy (while early stopping when the delta gets too large), we store two values in the NODE structure: *exp_idx* and *to_exp*, which store the index into the sorted list of expansions for this state, and what that state is, respectively. We keep nodes in the priority queue ordered from lowest to highest by their delta energy value, including the delta of the current expansion (indicated by *exp_idx*) they are on. When we process a node, we add a child node that is the result of applying that current expansion. We then reinsert the current node after advancing it to the next expansion (or not, if the delta is too high or we ran out of expansions). We can update the node’s energy value for the next expansion by subtracting the current expansion’s delta and adding the next one’s. Each child node stores the index of the expansion it used in the parent, *parent* _*exp_idx*, which is a snapshot of the parent’s *exp_idx* at the time the child was created.

Another difficulty is handling the extra unexpanded states when processing a node. If a particular state wants to make a multi-loop, that maps to two child states (the left and right side when decomposing the multi-loop). When we create a child, it only refers to a single state which it will continue to expand (*to_exp* in the NODE structure). But at some point we need to process the other state. This is handled by *unexp_idx* and *unexp_exp_idx* which record the next ancestor node which has an unexpanded state we need to eventually process, and the expansion that the unexpanded state is in, respectively. When we run out of expansions to process at a node, we start looking at the previously unexpanded states. We do this by checking if the node has an *unexp_idx*. If it does, we get the unexpanded state and create a new node to expand it. We also set the *unexp_idx* and *unexp_exp_idx* values of that new node to the unexpanded ancestor’s values. In this way, we maintain an implicit persistent linked list up through the tree’s ancestors with nodes that have unexpanded states. When processing a node with a new unexpanded state, we insert it into the tail of this linked list. When processing an unexpanded state, we delete it from the linked list by updating that node’s *unexp_idx*. Each node has different pointers to ancestors, so in totality *unexp_idx* works to maintain a tree of unexpanded states on top of the node tree, and each node processes the path from it to the root (the linked list) in this tree.

When we reach a node that has nothing to expand we return that as a complete suboptimal structure. Since we produce suboptimal structures one by one, this PFS can be terminated at any time, allowing us to iteratively sample the next suboptimal structure in order of energy. Unlike the iterative deepening approach, this does not require sometimes paying large time cost to get the next structure.

Note that it is imperative for performance and the correct asymptotic complexity to order the priority queue so that higher indexed nodes (nodes we found later) are popped before lower indexed nodes with the same energy. This creates a depth first order within the same energy value so evolving a partial structure to a complete suboptimal structure is prioritised. Without this, the algorithm could end up exploring every partial structure at a particular energy level (of which there appears to be an exponential amount) before producing any single structure.

In Figure 2 we show an example of how the PFS evolves a partial structure, and how the priority queue and persistent structures are updated as we traverse the expansion tree. In particular, the figure shows how the backlinks storing the implicit linked list of unexpanded states work.

**Figure 2:**
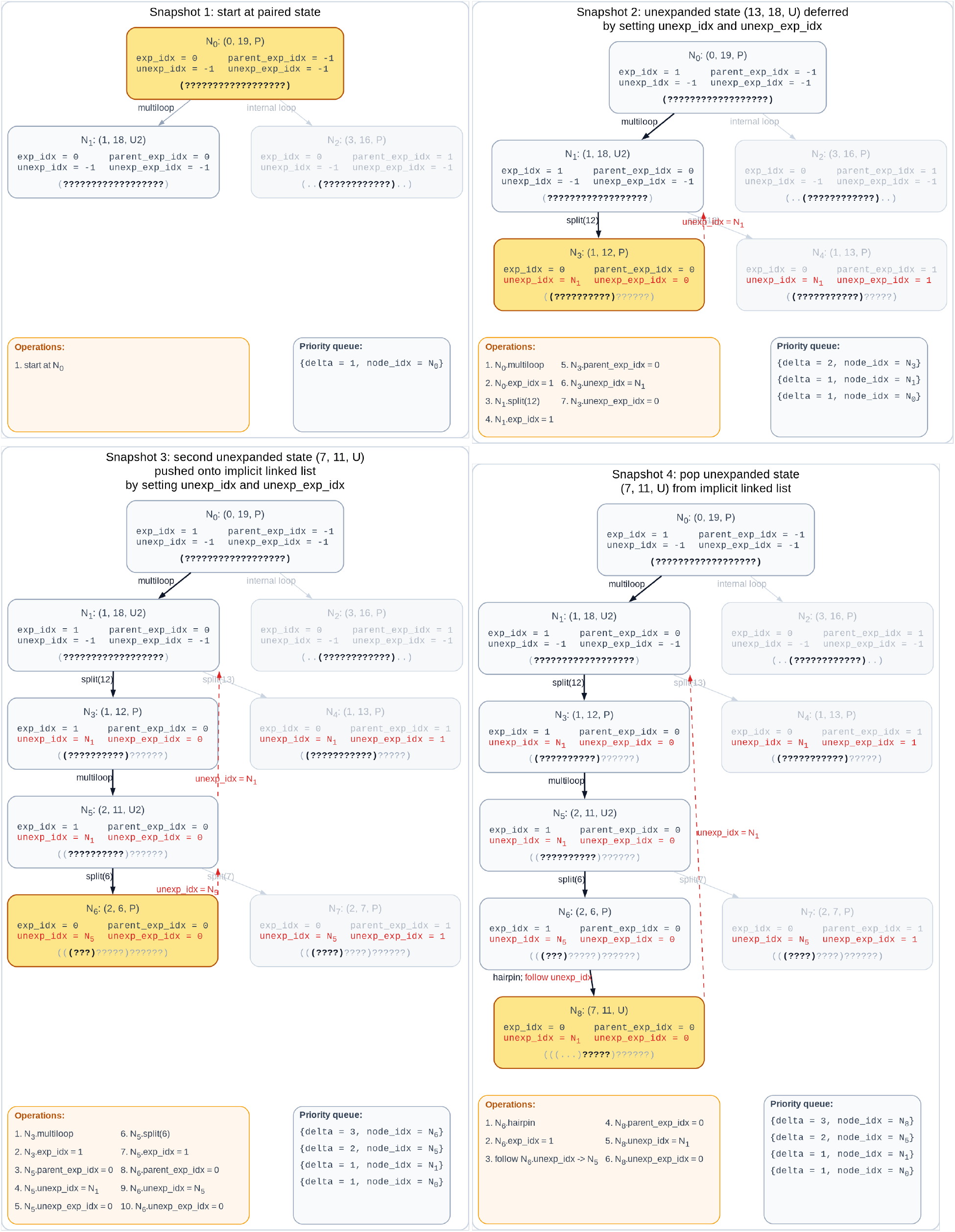
We illustrate the expansion tree and the evolution of a partial structure in the persistent suboptimal folding algorithm. Items shown in red are related to the handling of unexpanded states. Here we use the number of base pairs (Nussinov-like [30]) as the delta value, instead of difference to MFE, for simplicity.

### 2.3 Complexity analysis

We analyse the time and memory complexity in terms of the RNA length *N*, and the number of structures, *S*, or energy delta, *δ*.

In order to analyse the time and memory complexity in terms of *δ*, we make a simplifying (but supported) assumption about the distribution of number of suboptimal structures versus their energy values. It appears to form a normal distribution over the entire thermodynamic ensemble — for example, see Figure 3. This happens in both the *d2* mode (`ViennaRNA`’s default mode with approximate coaxial stacking) and all CTD modes (`RNAstructure`’s mode, including full coaxial stacking).

**Figure 3:**
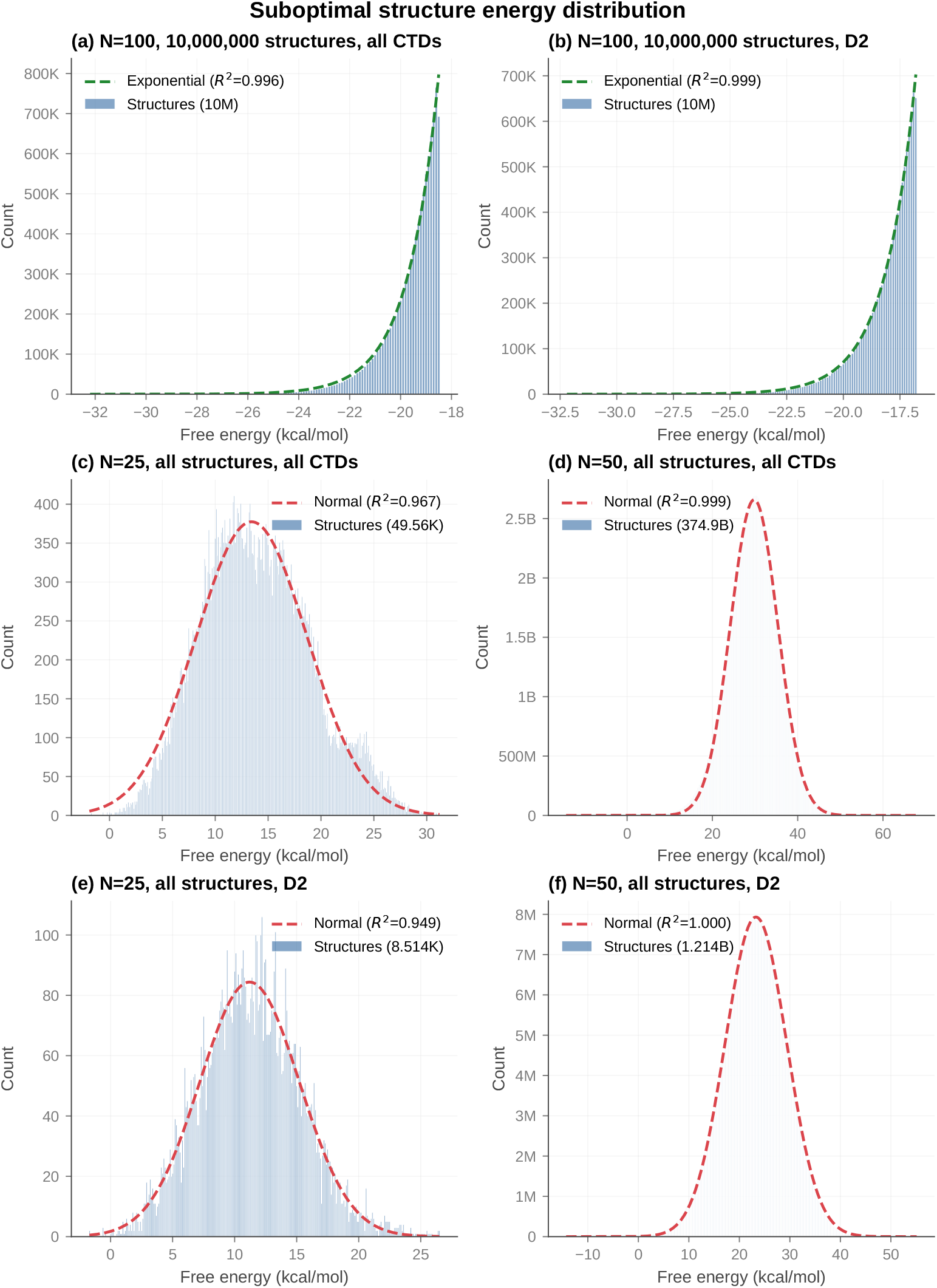
Energy distributions of sampled RNA structures across select RNA lengths (*N* = 25, 50, 100), showing a tendency towards a normal distribution in both `d2` mode and all CTDs mode. (a): The energy distribution of the first 10 000 000 structures with CTDs of a length 100 RNA sequence. It is fitted very well by an exponential function. (b): The energy distribution of the first 10 000 000 structures with `ViennaRNA d2` mode handling of CTDs of a length 100 RNA sequence. It is fitted very well by an exponential function. (c): The energy distribution of all suboptimal structures with CTDs of a length 25 RNA sequence. It is approximately normal. (d): The energy distribution of all suboptimal structures with CTDs of a length 50 RNA sequence. It is approximately normal. (e): The energy distribution of all suboptimal structures with `ViennaRNA d2` mode handling of CTDs of a length 25 RNA sequence. It is approximately normal. (f): The energy distribution of all suboptimal structures with `ViennaRNA d2` mode handling of CTDs of a length 50 RNA sequence. It is approximately normal.

We do not prove this rigorously here but provide an intuitive argument. Consider generating a random suboptimal structure as a process where each “sub-structure” is a random variable whose value is selected from a list of sub-structures it could be. If you consider a random structure’s free energy to be the sum of random variables *X*_0_, *X*_1_, …, *X*_*n*_

#### Algorithm 3

Pseudocode describing a step in suboptimal folding using persistent data structures.

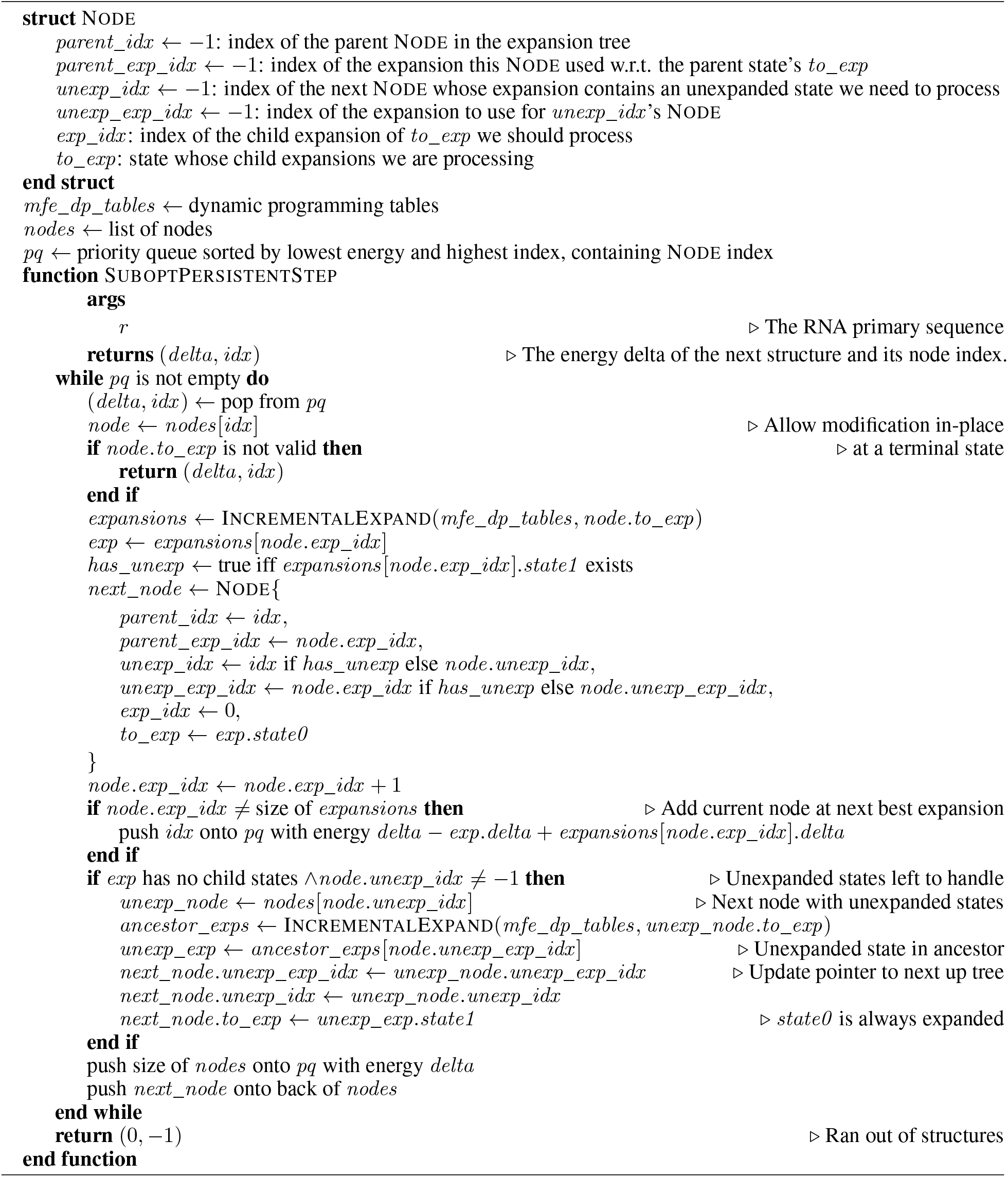

where each *X*_*i*_ represents the free energy of each random substructure, then application of the central limit theorem shows that random suboptimal structures should be normally distributed with respect to total free energy.

While the distribution over the entire thermodynamic ensemble is normal, when enumerating structures from lowest to highest energy, it starts on the left side which appears locally exponential. Generally speaking for large enough RNAs, the total number of suboptimal structures will be extremely large, so that most of structure generation will happen in this exponential regime. This is shown in Figure 3. The number of structures produced is also well approximated by *O*(*k*^*δ*^). This is important because it explains why iterative deepening is fast, as iterative deepening on an exponentially growing tree has the same time complexity as a normal DFS.

Similarly, we also make a simplifying assumption about the number of nodes in the expansion tree (*K*) with respect to the number of structures produced (*S*). We find that empirically, *K* = *O*(*S*) while in the exponential regime — see Figure 4. This is likely because each node in the expansion tree only looks at an *O*(1) number of children on average, in the exponential regime — that is, there are a lot of shared substructures. The following analysis uses this assumption, which we believe is well supported empirically. If this assumption is false, then *K* = *O*(*NS*) because the expansion tree cannot be larger than where each suboptimal structure produced has a completely disjoint path to the root, although this is not a tight bound.

**Figure 4:**
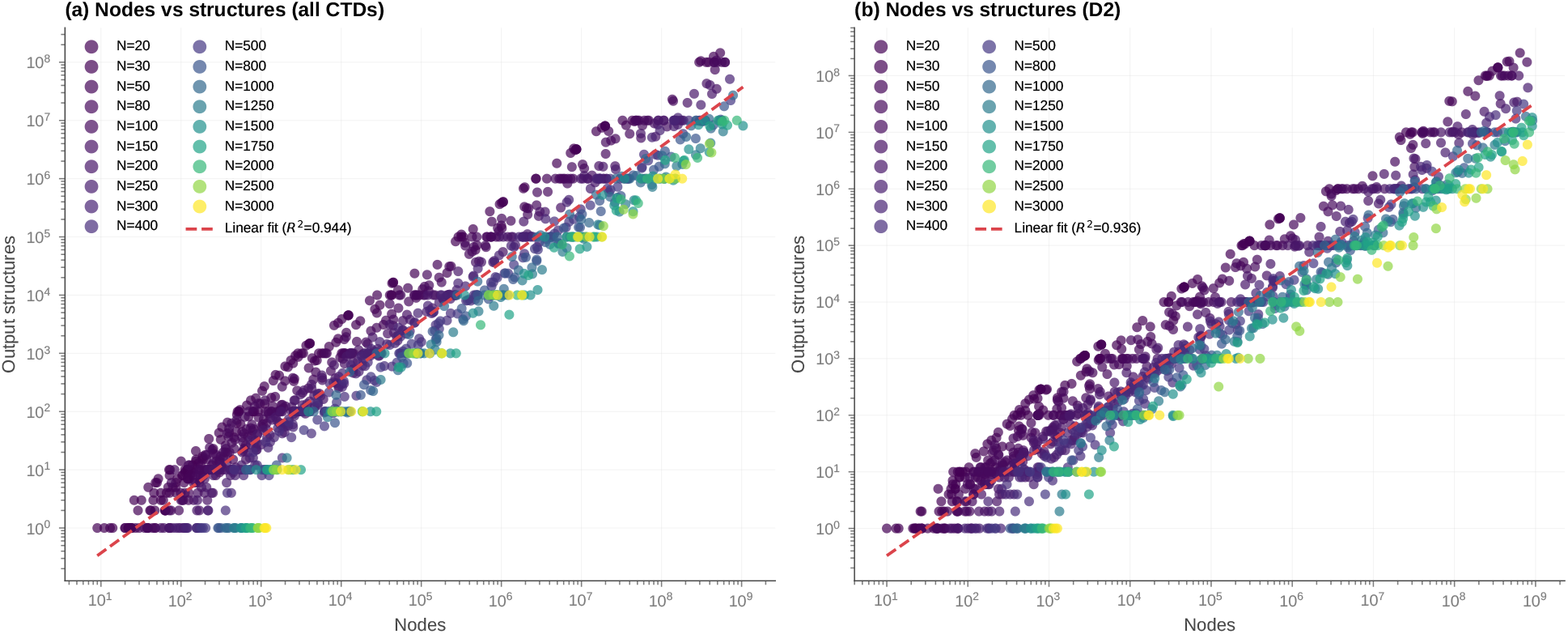
Expansion tree growth scales linearly with the number of generated structures under suboptimal folding regimes across various RNA lengths (*N*). (a): The distribution of number of nodes in the expansion tree vs the number of produced structures, for suboptimal folding with CTDs. It is approximately linear. (b): The distribution of number of nodes in the expansion tree vs the number of produced structures, for suboptimal folding with `d2` mode CTD handling. It is approximately linear.

We have summarised the time and memory complexity for each of the `memerna` algorithm variants in Table 4, the proofs of which are in Supplementary Section C.2. We write the complexities with the assumption that each structure will be copied when generated — with the persistent variant, generating the structure is a separate walk of the tree taking *O*(*N*) time, but if you do not need the full structure materialised then you would not need to pay the *O*(*N*) cost per structure. Based on our benchmarks, we have computed empirical complexities which corroborate the analysis here in Supplementary Figure A.

**Table 4:** Worst case complexity summary of `memerna` algorithm variants for sampling *S* structures. While the persistent variants have higher theoretical time complexity, the time to produce the next structure is limited to *O*(*N* log *S*), whereas the iterative deepening variants may need to re-run the DFS. The amortised complexity of the iterative deepening approach is better, however, highlighting the trade-off in predictability of marginal runtime. The amortised runtimes include *O*(*N*) work to copy the structure out of the global state for the iterative variant, and *O*(*N*) work to walk the tree for the persistent variant.

| Variant | Time (worst case) | Memory (worst case) | Time for one structure (worst case) | Time for one structure (amortised) |
| --- | --- | --- | --- | --- |
| iterative-strucs | $O(SN + N^3)$ | $O(N^3)$ | $O(SN)$ | $O(N)$ |
| iterative-delta | $O(k^\delta N + N^3)$ | $O(N^3)$ | $O(k^\delta N)$ | $O(N)$ |
| persistent-strucs | $O(S \log S + SN + N^3)$ | $O(S + N^3)$ | $O(N \log S)$ | $O(\log S + N)$ |
| persistent-delta | $O(\delta k^\delta \log k + k^\delta N + N^3)$ | $O(k^\delta + N^3)$ | $O(N \log S)$ | $O(\log S + N)$ |

## 3 Methods

### 3.1 Benchmark methodology

We ran benchmarks on a computer with a Ryzen 9950X3D CPU and 128 GiB of RAM. We gave each program a memory limit of 8 GiB and a time limit of 30 minutes per invocation.

We ran each test case of a particular length on ten randomly generated primary structures of lengths 100 and 1000 (generated once, so they were the same between invocations of different programs) and took the arithmetic mean with a bootstrapped 95% confidence interval around them, as shown in Figure 5. See Supplementary Table E for which RNA lengths and parameters we tested. See Supplementary Table D for how we generated the data and ran the benchmarks.

**Figure 5:**
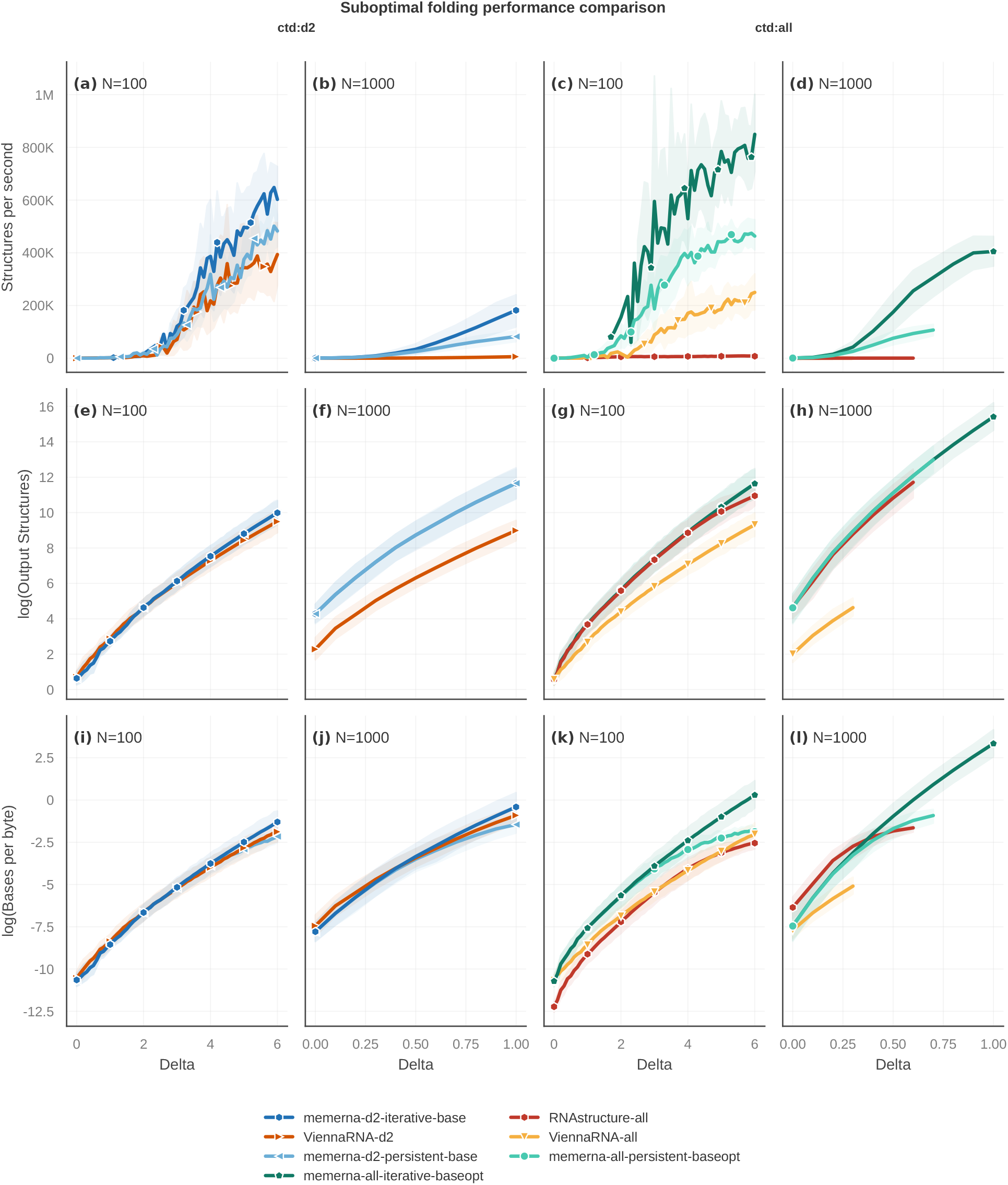
`memerna` iterative sampling benchmarking comparison. (a–b): Comparison of structures per second at differing RNA lengths in `d2` mode. (c–d): Comparison of structures per second at differing RNA lengths with all CTDs. (e–h): Comparison of the log of the number of output structures. (i–l): Comparison of the log of the number of bases produced per byte of maximum resident set size memory usage.

Different packages generate differing numbers of structures for the same input sequence and given delta. `ViennaRNA`, particularly for the length 1000 structures generated significantly fewer structures than `memerna`. To try to compare between packages we therefore look at structures per second and bases generated per byte of memory usage — that is, normalised to what each program actually output. We summarise the structure count ratios in Supplementary Table A.

We list the tested configurations in Supplementary Table B. See Supplementary Table C for the exact commands we ran for each program and configuration. We compare `memerna` using its `d2` mode with `ViennaRNA`, and `memerna` using its default mode of handling all CTDs with `RNAstructure`. This is the fairest mode of comparison, since `ViennaRNA` with its `d3` mode (which is approximately the same as RNAstructure’s CTD handling) appears less well supported.

## 4 Results

### 4.1 Iterative sampling outperforms existing suboptimal folding implementations

Across both energy models tested (dangles-only and full Turner 2004 with all CTDs), `memerna` consistently achieved the highest throughput measured as structures per second (Figure 5a–d). We observed this performance advantage at both 100 nt and 1000 nt sequence lengths, and it persisted as structure counts increased. In addition to higher throughput, `memerna` frequently enumerated a larger number of total structures within the same energy window, particularly for longer sequences (Figure 5e–h). We summarise the speedup over all data in Table 5.

**Table 5:**
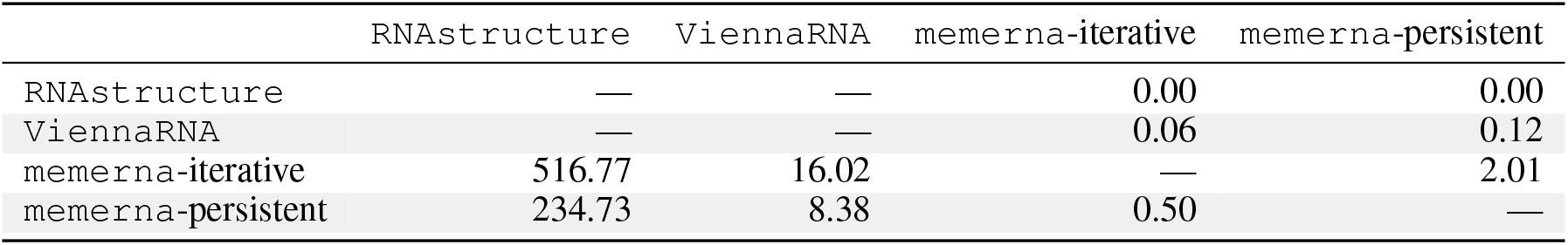
Pooled structures per second ratio. Values greater than 1 indicate that the row program is faster. For ViennaRNA we only included the d2 mode, and compared it to the d2 mode in memerna.

The iterative deepening implementation of `memerna` achieved memory efficiency comparable to `ViennaRNA` while sustaining higher structural output. In contrast, the persistent implementation exhibited increasing memory usage as structure counts grew, consistent with its retention of partial structures rather than depth-first traversal. At high structure counts, this led to greater peak memory consumption than `ViennaRNA`. We summarise the memory per base ratio over all data in Table 6.

**Table 6:** Pooled memory-per-base ratio. Values greater than 1 indicate that the row program uses more memory per base. For ViennaRNA we only included the d2 mode, and compared it to the d2 mode in memerna.

|  | RNAstructure | ViennaRNA | memerna-iterative | memerna-persistent |
| --- | --- | --- | --- | --- |
| RNAstructure | — | — | 5.20 | 1.34 |
| ViennaRNA | — | — | 2.23 | 0.14 |
| memerna-iterative | 0.19 | 0.45 | — | 0.01 |
| memerna-persistent | 0.75 | 6.93 | 78.95 | — |

These results demonstrate that `memerna`’s iterative deepening algorithm provides superior enumeration throughput while maintaining memory efficiency. These results establish iterative sampling as an efficient alternative to existing suboptimal folding approaches, motivating its application to experimentally guided RNA folding analyses.

### 4.2 Iterative sampling improves efficiency and data consistency of sampled structures in R2D2

`R2D2` reconstructs out-of-equilibrium RNA folding intermediates using a sample-and-select framework, where candidate RNA secondary structures are first generated and evaluated against experimental probing data using a distance metric *D*, selecting *D*_min_ structure (Figure 6a, Supplementary Section D.2) [4]. In its original implementation, candidate structures are generated via stochastic Boltzmann sampling, which inherently biases sampling towards low-free-energy structures. This bias, combined with stochasticity, leads to redundant sampling and inefficient exploration of alternative, high-free-energy structures that may better align with what is captured experimentally.

**Figure 6:**
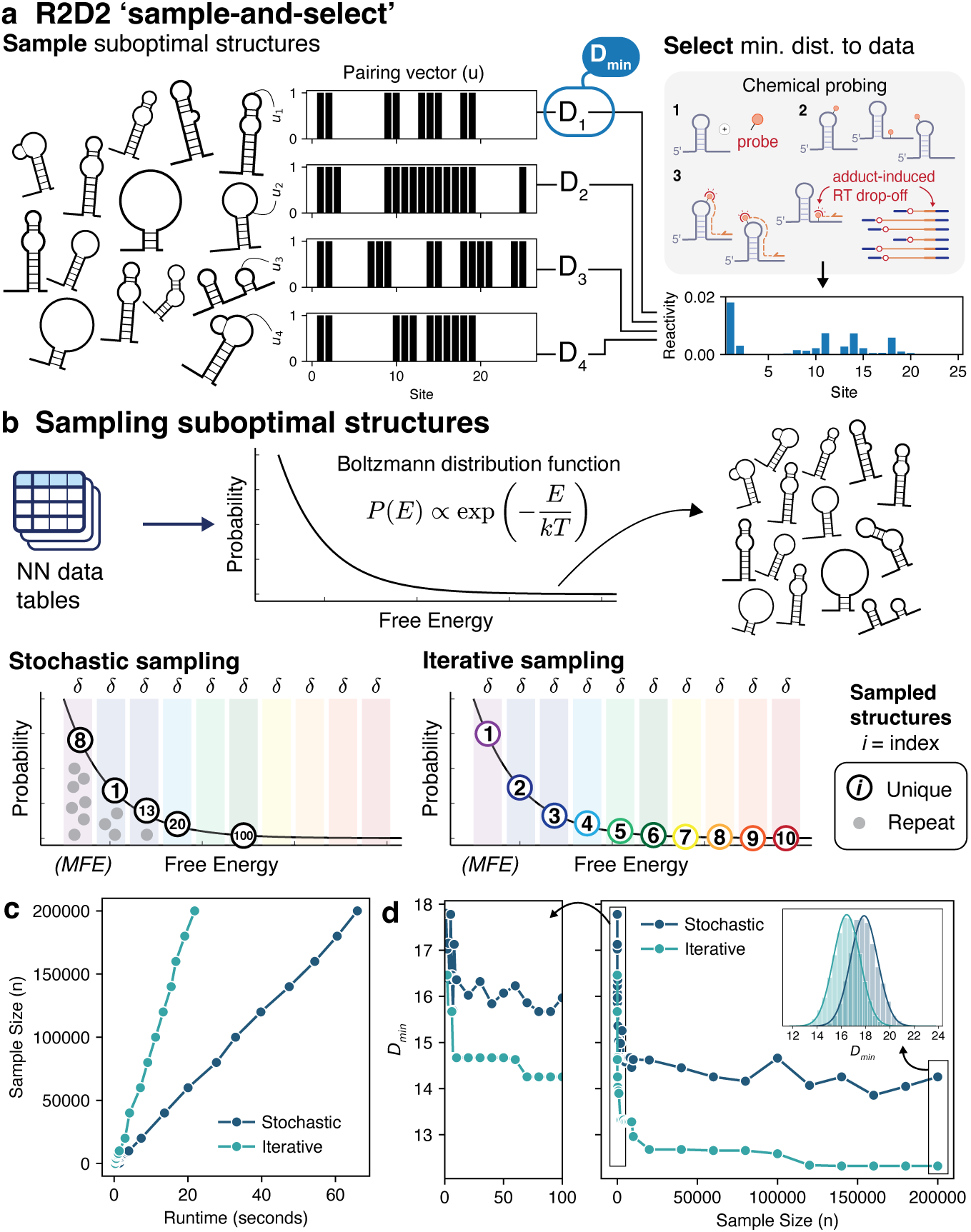
Iterative sampling in `memerna` results in immediate efficiency and performance gains when applied to `R2D2`’s sample-and-select scheme (a): `R2D2` sample-and-select scheme for identifying out-of-equilibrium structures. A sampling algorithm generates suboptimal structures, and a parametric distance function, *D*, quantifies agreement between each sampled secondary structure and experimental structure probing data. The structure with the minimum distance *D* is then selected. (b, top): Suboptimal structure sampling using nearest-neighbour (NN) thermodynamic models produces a Boltzmann weighted ensemble. Structures are sampled on this energy landscape. (b, bottom): Comparison of stochastic sampling and `memerna`’s iterative sampling over a hypothetical energy landscape. `memerna` iterative sampling was applied to the *E. coli* SRP RNA 111 nt cotranscriptional SHAPE-Seq dataset. (c): Runtime versus number of sampled structures when using iterative (`memerna`) and stochastic (`RNAstructure`) sampling approaches, demonstrating improved computational efficiency. (d): Increasing sample size with `memerna` yields structures with lower *D* values than `RNAstructure`. Inset: Histograms of *D*, coloured as in the main plots, show a clear shift toward better-fitting (lower *D*) structures.

To address this limitation, we replaced stochastic sampling with deterministic iterative sampling implemented in `memerna` (Algorithm E). Unlike stochastic approaches, iterative sampling enumerates unique structures in order of increasing free energy (Figure 6b, Supplementary Figure Bb), completely eliminating redundancy while ensuring uniqueness of each sampled structure.

Applying this approach to cotranscriptional SHAPE-seq data for *E. coli* SRP RNA (111 nt) yields immediate gains in both efficiency and performance (Figure 6c–d). Runtime scaling shows that iterative sampling generates substantially more structures within the same time (≈ 200,000 vs. ≈50,000 structures in 20 s; Figure 6c). Moreover, the minimum *D* decreases more rapidly and reaches a lower final value as sampling depth increases, indicating improved identification of data-consistent structures (Figure 6d). Histograms of *D* further reveal a systematic shift of the entire sampled distribution toward better-fitting structures (lower *D*) (Figure 6d, inset), indicating enhanced agreement with experimental data across the sampled ensemble.

These results demonstrate that deterministic iterative sampling improves `R2D2` as a drop-in replacement by removing redundant sampling and enabling more efficient search for data-consistent structures, a feature that extends to other sampling-based approaches [2]. Moreover, because sampling is deterministic and non-redundant, structures can be enumerated until a convergence criterion (e.g., stabilisation of *D*_min_) is reached, providing a practical stopping condition not available with stochastic sampling.

### 4.3 Probing-guided sampling reshapes the folding landscape

Since the development of partition function algorithms [27], NN models have enabled RNA ensemble generation. However, Boltzmann bias and stochastic sampling limit their ability to capture suboptimal structures comprehensively, a key limitation for RNAs that fold out of equilibrium. Iterative sampling overcomes this by enumerating suboptimal structures in order of energy, substantially expanding the accessible conformational space (Supplementary Section C.3). This enables direct interrogation of how entire ensembles respond to perturbations. Here, we examined the impact of chemical probing data, which improves minimum free energy (MFE) structure prediction accuracy by introducing a pseudo-free energy term that perturbs NN parameters [31]. We hypothesised that the resulting restrained energy model would translate to a shifted energy landscape, producing a distinct set of suboptimal structures (Figure 7b).

**Figure 7:**
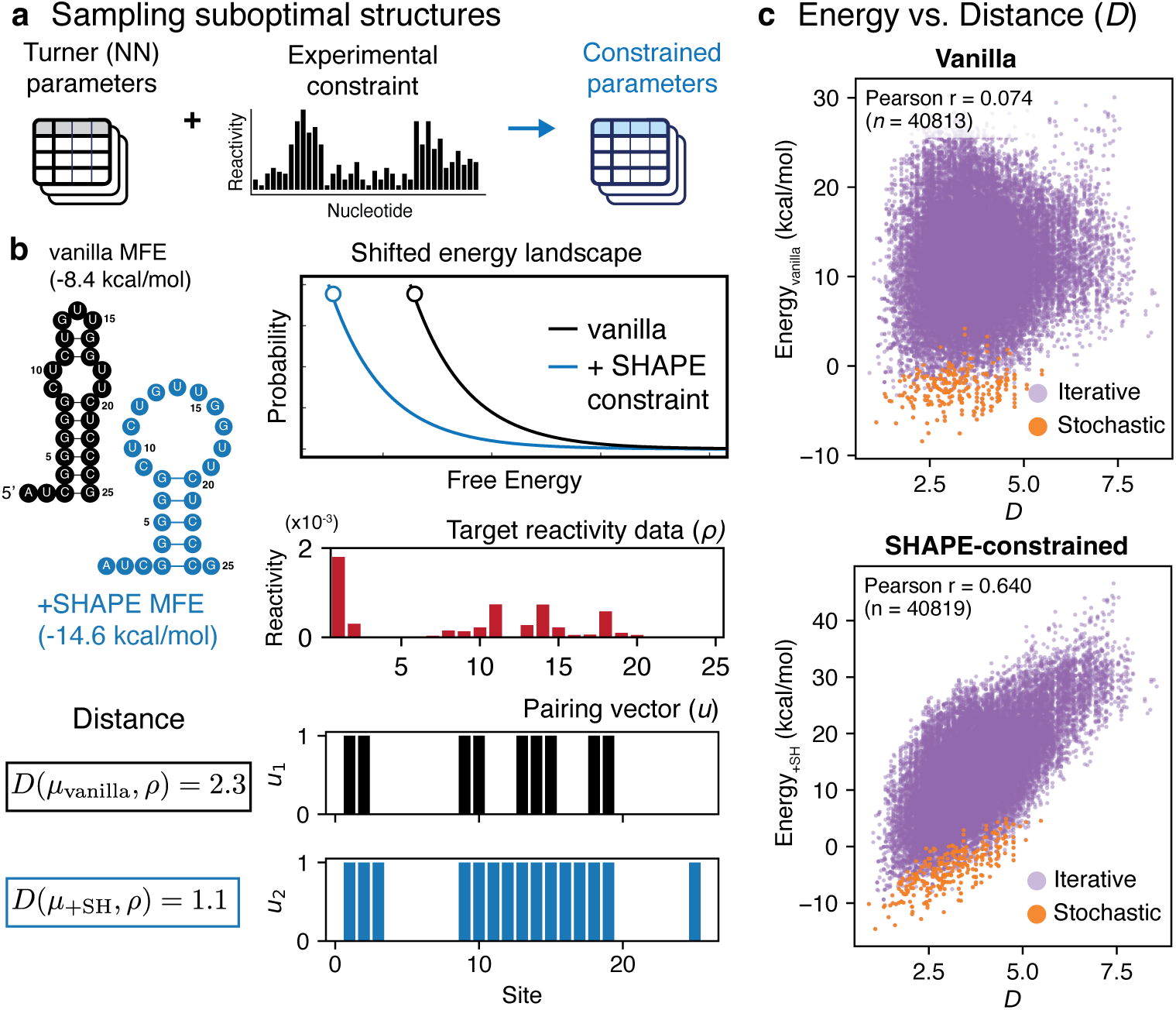
Restraining sampling with chemical probing data reshapes the sampled free energy landscape. (a): Turner nearest-neighbour parameters can be modified by incorporating experimental chemical probing data as pseudo-free energy penalties. (b): These restraints reshape the underlying energy landscape, leading to sampling of structures that differ from those obtained under unrestrained (“vanilla”) conditions. MFE structures shown for “vanilla” and SHAPE-restrained (“+SHAPE”) sampling. Experimental chemical probing reactivity data (*ρ*) for the 25-nt *E. coli* SRP RNA (red) are shown alongside the binary pairing vectors corresponding to the vanilla MFE structure, *u*_vanilla_ (black) and the SHAPE-restrained MFE structure, *u*_+SH_ (blue). The distance values between each structure and the experimental data are *D*(*u*_vanilla_, *ρ*) = 2.3 and *D*(*u*_+SH_, *ρ*) = 1.1, respectively. (c): Left: Free energy versus *D* for 50,000 sampled structures under vanilla (unrestrained) sampling. Structures also recovered by stochastic sampling are highlighted in orange. Free energy shows weak correlation (*r* = 0.074) with agreement to SHAPE data. Right: The same analysis with SHAPE restraints applied during sampling. Restraints reshape the energy landscape, producing a strong positive correlation between free energy and *D* (*r* = 0.64), thus biasing sampling towards generating structures that are consistent with experimental probing data.

We first applied iterative sampling to a control model system – a 25-nt sequence at the 5^*′*^ end of the *E. coli* SRP RNA (Supplementary Figure Ba) [32], [33], expected to form a hairpin. Under standard NN parameters, the minimum free energy (MFE) structure is a hairpin containing an internal loop (-8.4 kcal/mol). In contrast, incorporating SHAPE restraints yields a distinct structure with a shortened stem and enlarged loop (-14.6 kcal/mol), representing a ≈ 6 kcal/mol stabilisation. Consistent with this shift, the `R2D2` distance metric *D* indicates that the SHAPE-derived MFE is more consistent with experimental data. To generalise this effect to the entire sampled ensemble, we quantified the relationship between free energy and *D* across sampled ensembles. Under unrestrained (vanilla) sampling, the free energy of sampled structures shows little correlation with *D* (Pearson *r* = 0.074) (Figure 7c). In contrast, restrained sampling produces a strong correlation (*r* = 0.640), such that lower-energy structures are also those most consistent with the experimental data (Figure 7c). In this system, the lowest-energy structure is also the lowest-*D* structure and is recovered first by the iterative algorithm. More broadly, structures emerge in a predictable order: the dominant, data-consistent fold appears first, followed by higher-energy conformations that progressively deviate in *D* from the experimental profile.

Together, these results indicate that incorporating chemical probing reactivities within iterative sampling reweights NN parameters such that free energy becomes aligned with experimental consistency, effectively transforming Δ*G* into a proxy for experimental likelihood and obviating the need for heuristic distance-based selection.

### 4.4 Distinct ensembles emerge under equilibrium and cotranscriptional folding conditions

Recent advances in chemical probing techniques enable direct measurement of out-of-equilibrium (OOE) cotranscriptional folding (Figure 8a) [4], [17], [21]. A central challenge is to infer the underlying structures from these data, as NN models are inherently thermodynamic and do not account for kinetic effects. The original `R2D2` framework addressed this by promoting structural diversity through restrained stochastic sampling and distance-based selection (Figure 6a); however, this approach required parameter tuning and sequence-dependent optimisation of runs. In contrast, iterative sampling enables exploring how restrained energy landscapes generate distinct structural ensembles, enabling direct and deterministic comparison of folding landscapes across conditions.

**Figure 8:**
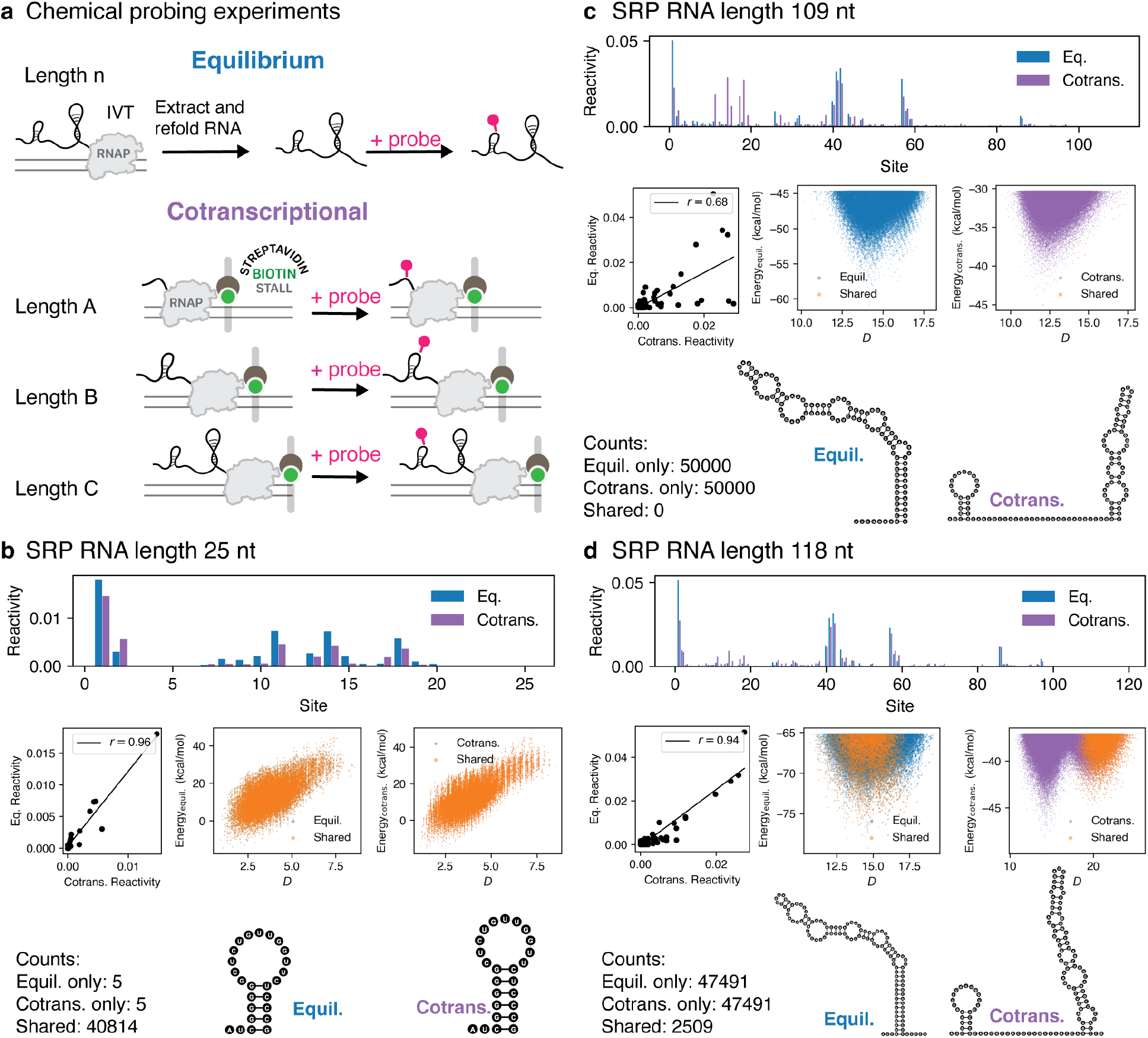
Comparison of equilibrium vs. cotranscriptional folding landscapes using SHAPE-restrained iterative sampling. (a): Schematic illustration of the difference between equilibrium refolding and cotranscriptional RNA chemical probing experiments. In equilibrium refolding, RNAs are generated by transcription, extracted and refolded in equilibrium conditions before probing. In cotranscriptional folding, stalled transcription elongation complexes are used to probe RNAs that are emerging from the RNA polymerase. (b): Analysis of the 25-nt *E. coli* SRP RNA under equilibrium and cotranscriptional conditions, organised in three rows. (top row) Bar plots of nucleotide reactivities measured under equilibrium and cotranscriptional conditions. (middle row) Left, Pearson correlation (*r*) between the two reactivity profiles. Middle and right, free energy versus *D* for structures sampled with SHAPE restraints derived from equilibrium data (blue) or cotranscriptional data (purple). Structures shared between the two ensembles are highlighted in orange (“shared”). (bottom row) Centroid structures for equilibrium and cotranscriptional ensembles, annotated with the number of structures unique to each condition and the number shared between conditions. (c): As in (b) but for length 109 nt. (d): As in (b) but for length 118 nt.

We applied SHAPE-restrained iterative sampling across selected transcript lengths of the SRP RNA probed to track how ensemble evolves under equilibrium and cotranscriptional conditions. At 25 nt, highly correlated reactivity profiles yield nearly identical structural, indicates that early folding is dominated by a shared, near-equilibrium structure (Figure 8b). In contrast, by 109 nt, the two conditions diverge markedly (Pearson *r* = 0.68), and iterative sampling reveals distinct basins between the two conditions occupying distinct energy regimes (Figure 8c). Although the ensembles appear to abruptly cut off at around 45 kcal/mol, consistent with incomplete sampling, the two ensembles nevertheless remain fully separated. The cotranscriptional ensemble is dominated by a kinetically trapped architecture consisting of a short 5^*′*^ hairpin (H1, 5 ≈ bp) and a truncated 3^*′*^ hairpin (≈ 16 bp), whereas the equilibrium ensemble favours an elongated ≈ 32-bp hairpin. This divergence recapitulates prior `R2D2` findings, but here emerges directly from the full sampled landscape rather than distance-based selection [4].

By 118 nt, partial convergence re-emerges as reactivity profiles become more similar and shared structural classes reappear (Figure 8d). However, the ensembles remain heterogeneous, with equilibrium states only partially represented in the cotranscriptional ensemble, while a cotranscription-specific low *D* basin persists (Figure 8d). Its centroid (Supplementary Section D.6) resembles the kinetically trapped architecture observed at 109 nt, suggesting a gradual transition rather than an abrupt switch. Notably, this competing conformational basin was not resolved by prior `R2D2` distance-minimisation analysis, which collapses the ensemble to a single structure.

Importantly, these results demonstrate that correlation between probing datasets alone does not capture differences in ensemble organisation. Although 25-nt (Pearson *r* = 0.96) and 118-nt (Pearson *r* = 0.94) datasets appear similarly correlated, their sampled landscapes differ fundamentally: the former shows complete structural overlap, whereas the latter exhibits energetic re-weighting, partial overlap, and condition-specific conformational basins.

Together, these results show that SHAPE-restrained iterative sampling not only reproduces known folding transitions but also has the power to reveal nuanced aspects of how experimental conditions reshape structural ensembles that are hidden by distance-based sample-and-select approaches. Moreover, by enabling sampling to convergence, iterative sampling provides a principled way to recover equilibrium folds while retaining intermediate states. While these analyses reveal qualitative differences between folding landscapes, a quantitative framework is required to systematically measure their divergence.

### 4.5 Ensemble divergence reveals cotranscriptional folding dynamics

Building on these qualitative differences, we next quantified divergence between equilibrium and cotranscriptional ensembles quantitatively at the level of structural probability distributions (Figure 8). For each sampled structure, *s*, its probability (Equation (A)) is determined by reactivity-restrained Δ*G*_*s*_ and the corresponding partition function *Q*, such that the same structures may have distinct probabilities under different folding regimes (Figure 9a). To quantify the difference between cotranscriptionally-restrained and equilibrium-restrained distributions, we applied the Jensen-Shannon divergence (JSD) to their structural probability distributions, where values of 0 indicate identical ensembles and 1 indicates completely non-overlapping ensembles (Figure 9a, Supplementary Section D.7).

**Figure 9:**
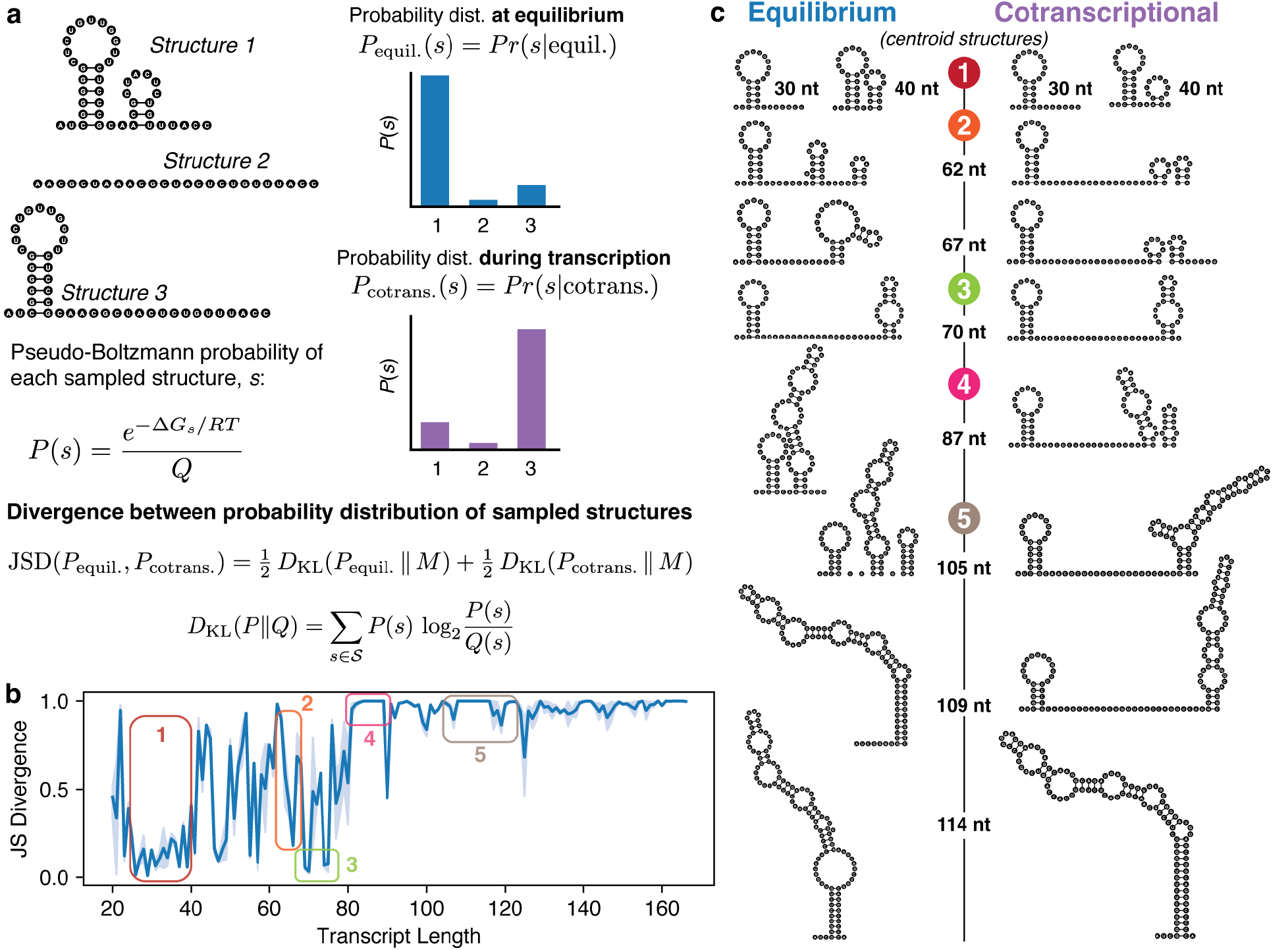
Quantifying divergence between structural probability distributions generated from equilibrium and cotranscriptional probing data across RNA lengths. (a): Jensen-Shannon divergence (JSD) is used to quantify differences between structural probability distributions derived from free energies (Δ*G*_*s*_). Structure probabilities (*P* (*s*)) are computed from Δ*G*_*s*_ values using partition functions (*Q*) calculated with nearest-neighbour (NN) energy models. Incorporation of probing restraints modifies Δ*G*_*s*_ and *Q*, thereby reshaping the structural ensemble. JSD is computed using the average distribution 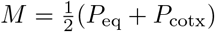, where *P*_eq_ and *P*_cotx_ denote the equilibrium and cotranscriptional structural probability distributions, respectively. In the illustrative example, three candidate structures exhibit distinct equilibrium and cotranscriptional probabilities; JSD quantifies the divergence between these distributions. (b): JSD across SRP RNA transcript lengths comparing equilibrium refolding and cotranscriptional folding. Shaded regions indicate standard error across 2-3 cotranscriptional replicates. Transcript regions referenced in the text are highlighted by colour. (c): Centroid structures derived from the equilibrium (left) and cotranscriptional (right) ensembles at representative transcript lengths within the annotated regions (colours correspond to panel (b)).

Applying this framework to SRP RNA across transcript lengths (25-166 nt) reveals a dynamic trajectory of ensemble divergence (Figure 9b). At short lengths (<40 nt), JSD is near zero (Figure 9b, 1), indicating near identical structural ensembles. As transcription proceeds, JSD fluctuates (Figure 9b, 2), reflecting transient intermediate states without a dominant fold (Figure 9b, 2). Spikes in JSD correspond to cotranscriptional-specific intermediates, whereas dips indicate partial relaxation toward equilibrium-like conformations. Notably, a pronounced dip at 70 nt coincides with a putative RNAP pause site at (U82/U84), consistent with pause-enabled equilibration (Figure 9b, 3) [16]. Beyond ≈ 90 nt, JSD rises to sustained high values, indicating persistent divergence and largely non-overlapping ensembles (Figure 9b, 4-5).

To interpret the structural basis of this divergence, we analysed ensemble centroids across transcript lengths (Supplementary Section D.6). Cotranscriptional folding preferentially resolves newly transcribed sequence into local 3^*′*^ hairpins rather than reorganising upstream helices. From ≈ 60 to 113 nt, nascent 3^*′*^ segments consistently form short stem-loops(Figure 9c, regions 2-5), with 70 nt again marking an exception due to equilibration (Figure 9c, region 3). In contrast, equilibrium ensembles at the same lengths more readily sample globally rearranged conformations that incorporate upstream pairing partners, including the restructuring of the early-formed 5^*′*^ hairpin. These observations support a model in which rapid local folding during transcription stabilises upstream structures and delays global rearrangements first reported in [4], [made more accessible, deterministic and detailed by iterative sampling - need to add clause on what’s newly enabled]

This mechanism explains the timing of the tri-helix-to-elongated hairpin rearrangement of SRP [4]. Under cotranscriptional conditions, stabilisation of the 5^*′*^ hairpin delays formation of the tri-helical intermediate and the final elongated fold. While tri-helix formation is detected at ≈105 nt, stable adoption occurs only by ≈114 nt, consistent with optical tweezer measurements indicating cessation of “hopping” or commitment to the functional fold [34]. In contrast, the elongated conformation is energetically accessible under equilibrium refolding conditions by ≈ 94 nt, indicating that energetic feasibility precedes kinetic realisation.

Together, this analysis demonstrates that treating iteratively sampled structures as probabilistic ensembles enables direct, quantitative comparison of folding landscapes across conditions. Applied to SRP, JSD reveals kinetic trapping, pause-associated equilibration, and delayed structural commitment, providing a principled, deterministic, and interpretable framework for dissecting nonequilibrium folding dynamics along transcriptional trajectories.

## 5 Discussion

We present new iterative sampling algorithms including an iterative deepening method and a persistent data structure method. These improve upon existing algorithms by allowing a natural “iterator-like” sampling method. They also have excellent theoretical and empirical performance compared to existing methods. Both of these qualities are applied to great effect in `R2D2`, where they are an ideal fit. `R2D2` needs highly efficient sampling algorithms that are non-redundant and can handle arbitrary stopping criteria. We can sample many more structures than before. We were also able to sample the top 150,000 structures, enabling a fair comparison between the equilibrium ensemble and the chemical probing guided ensemble via pseudo-free energies. It should be observed that this would not easily be possible with existing algorithms. We have already pointed out that stochastic sampling is redundant. Also, using Wuchty’s algorithm is poorly defined with pseudo-free energies and it would be hard to pick an energy window that allows a fair comparison between equilibrium and pseudo-free energy guided samples.

We benchmarked our new algorithms against `ViennaRNA` and `RNAstructure` because they represent the most widely used and practical baseline for stochastic Boltzmann sampling in RNA secondary structure analysis. Although Wuchty-style enumeration reduces redundancy within a fixed energy window, it does not operate as a true iterator and does not include the optimisations introduced here. In contrast, the new deterministic algorithms implemented in `memerna` enumerate structures in strict free energy order, are non-redundant by construction, and function as genuine iterators. They also demonstrate excellent theoretical and empirical performance, even when considered purely as a sampling algorithm independent of these additional features. This enables scalable exploration of longer RNAs within a defined computational budget. At the same time, the framework inherits the standard limitations of nearest-neighbour secondary structure models, including the exclusion of pseudoknots and other tertiary effects.

Deterministic iterative sampling transforms ensemble analysis from probabilistic approximation to ordered landscape traversal. By eliminating the redundancy inherent to Boltzmann sampling, it enables systematic and reproducible exploration of condition-specific pseudoenergy landscapes defined by SHAPE-restrained Δ*G* values. In this work, we treat sampled structures directly as data. Structural ensembles become analysable datasets, allowing probability distributions over structures to be compared quantitatively, for example through Jensen–Shannon divergence. This shifts interpretation away from selecting a single best-fitting structure toward ensemble-level inference and establishes a computational framework that couples experimental restraints with deterministic structural ensembles. While SHAPE pseudo-free energies remain parametric and tunable, iterative sampling makes explicit how these restraints re-weight conformational landscapes and clarifies the need for more principled energetic formalisms in the future.

Finally, deterministic sampling provides a practical route to studying nonequilibrium folding. Identifying kinetic traps has traditionally required extensive experimental inference while computational approaches have been obscured by Boltzmann bias, which favours equilibrium-dominant structures. By expanding structural coverage in a non-redundant and reproducible manner, iterative sampling enables direct interrogation of out-of-equilibrium ensembles and complements existing cotranscriptional probing approaches. Applied to SRP RNA, this framework not only recapitulates the well-characterised tri-helix rearrangement but also reveals broader principles of cotranscriptional folding as a coupled and dynamically reweighted landscape. Together, these advances establish both a computational innovation in ordered secondary structure enumeration and a new analytical framework for translating experimental probing data into ensemble-resolved, deterministic quantities.

## 6 Source code

The code used for `R2D2` v2 is available on GitHub at https://github.com/LucksLab/r2d2_v2 and is archived at https://doi.org/10.5281/zenodo.19169630. Instructions for configuring and running the software are provided in the `README.md` file within the repository.

The `memerna` source code is available at https://github.com/Edgeworth/memerna/releases/tag/release-tag/0.2.2 and is archived at https://doi.org/10.5281/zenodo.19651365. Instructions for running `memerna` are in the `README.md` file in the repository.

## 8 Acknowledgments

We thank Jim Brink and Steve Hockema (496code) for developing and running `R2D2` v2. We thank Ryan Krueger for initial discussions and facilitating the collaboration between MW and JBL.

