## Supplementary Materials for "Efficient and scalable modelling of cotranscriptional RNA folding with deterministic and iterative RNA structure sampling"

### A Supplementary Figures

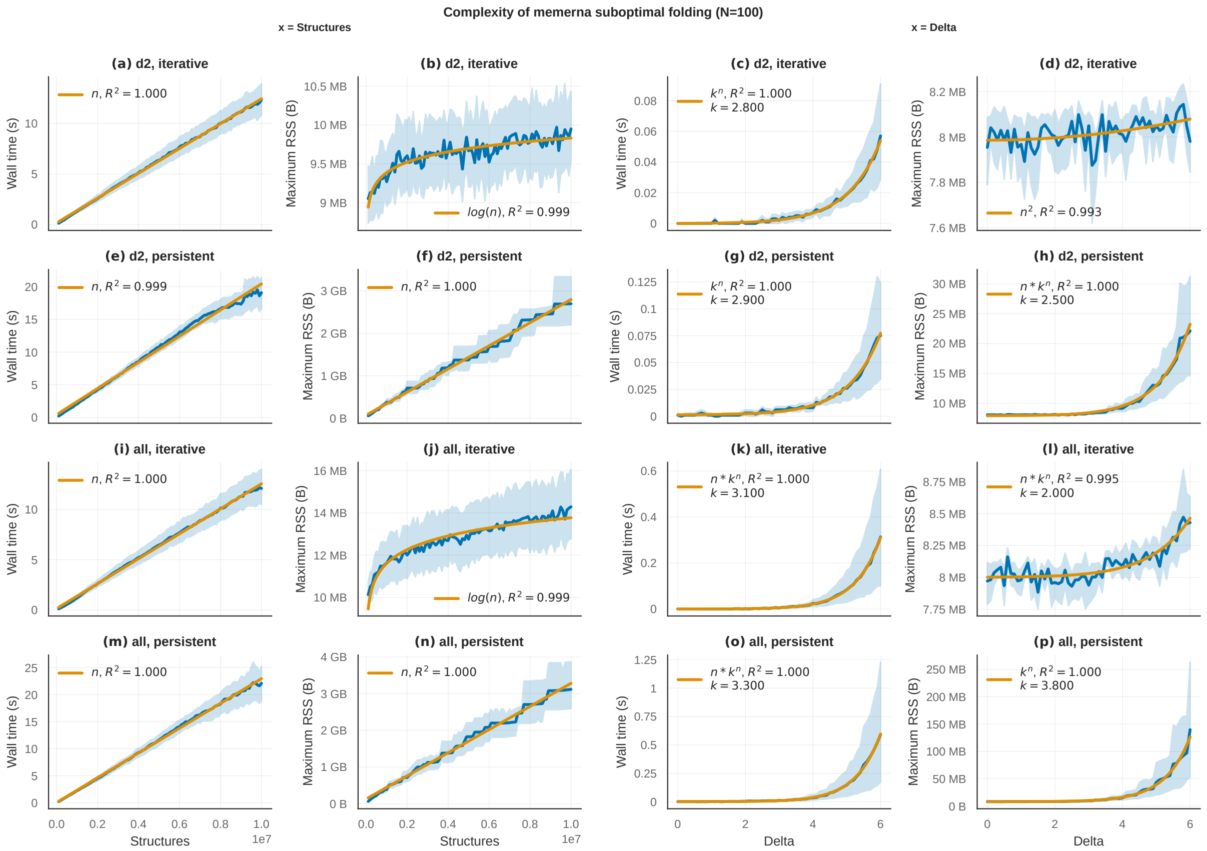

Figure A: Benchmark data with empirical complexities for `memerna` at $N = 100$. The left two columns plot wall time against the number of structures, and the right two columns plot wall time against delta. We selected the best fitting curve out of a selection of common functions. Some curves do not match the theoretical complexity analysis, but this is not because the theoretical complexity analysis is wrong, but rather that some data happens to be slightly better fit by a more complex function. (a): `d2` iterative, wall time by structures: $O(N)$, $R^2 = 1.000$. This matches the theoretical analysis. (b): `d2` iterative, memory by structures: $O(\log N)$, $R^2 = 0.999$. The theoretical analysis shows this is $O(1)$. The range of memory use is very small, so the empirical analysis is not correct here. (c): `d2` iterative, wall time by delta: $O(k^N)$, $k = 2.8$, $R^2 = 1.000$. This matches the theoretical analysis. (d): `d2` iterative, memory by delta: $O(N^2)$, $R^2 = 0.993$. This again occupies a very small memory range, so the theoretical analysis of $O(1)$ is correct. (e): `d2` persistent, wall time by structures: $O(N)$, $R^2 = 0.999$. This matches the theoretical analysis. (f): `d2` persistent, memory by structures: $O(N)$, $R^2 = 1.000$. This matches the theoretical analysis. (g): `d2` persistent, wall time by delta: $O(k^N)$, $k = 2.9$, $R^2 = 1.000$. This matches the theoretical analysis. (h): `d2` persistent, memory by delta: $O(N \cdot k^N)$, $k = 2.5$, $R^2 = 1.000$. This mostly matches the theoretical analysis, but includes an extra factor of delta, which is a small value compared to the noise. (i): all CTDs iterative, wall time by structures: $O(N)$, $R^2 = 1.000$. This matches the theoretical analysis. (j): all CTDs iterative, memory by structures: $O(\log N)$, $R^2 = 0.999$. The theoretical analysis shows this is $O(1)$. The range of memory use is very small, so the empirical analysis is not correct here. (k): all CTDs iterative, wall time by delta: $O(N \cdot k^N)$, $k = 3.1$, $R^2 = 1.000$. This mostly matches the theoretical analysis. (l): all CTDs iterative, memory by delta: $O(N \cdot k^N)$, $k = 2.0$, $R^2 = 0.995$. This mostly matches the theoretical analysis. (m): all CTDs persistent, wall time by structures: $O(N)$, $R^2 = 1.000$. This matches the theoretical analysis. (n): all CTDs persistent, memory by structures: $O(N)$, $R^2 = 1.000$. This matches the theoretical analysis. (o): all CTDs persistent, wall time by delta: $O(N \cdot k^N)$, $k = 3.3$, $R^2 = 1.000$. This mostly matches the theoretical analysis. (p): all CTDs persistent, memory by delta: $O(k^N)$, $k = 3.8$, $R^2 = 1.000$. This matches the theoretical analysis.

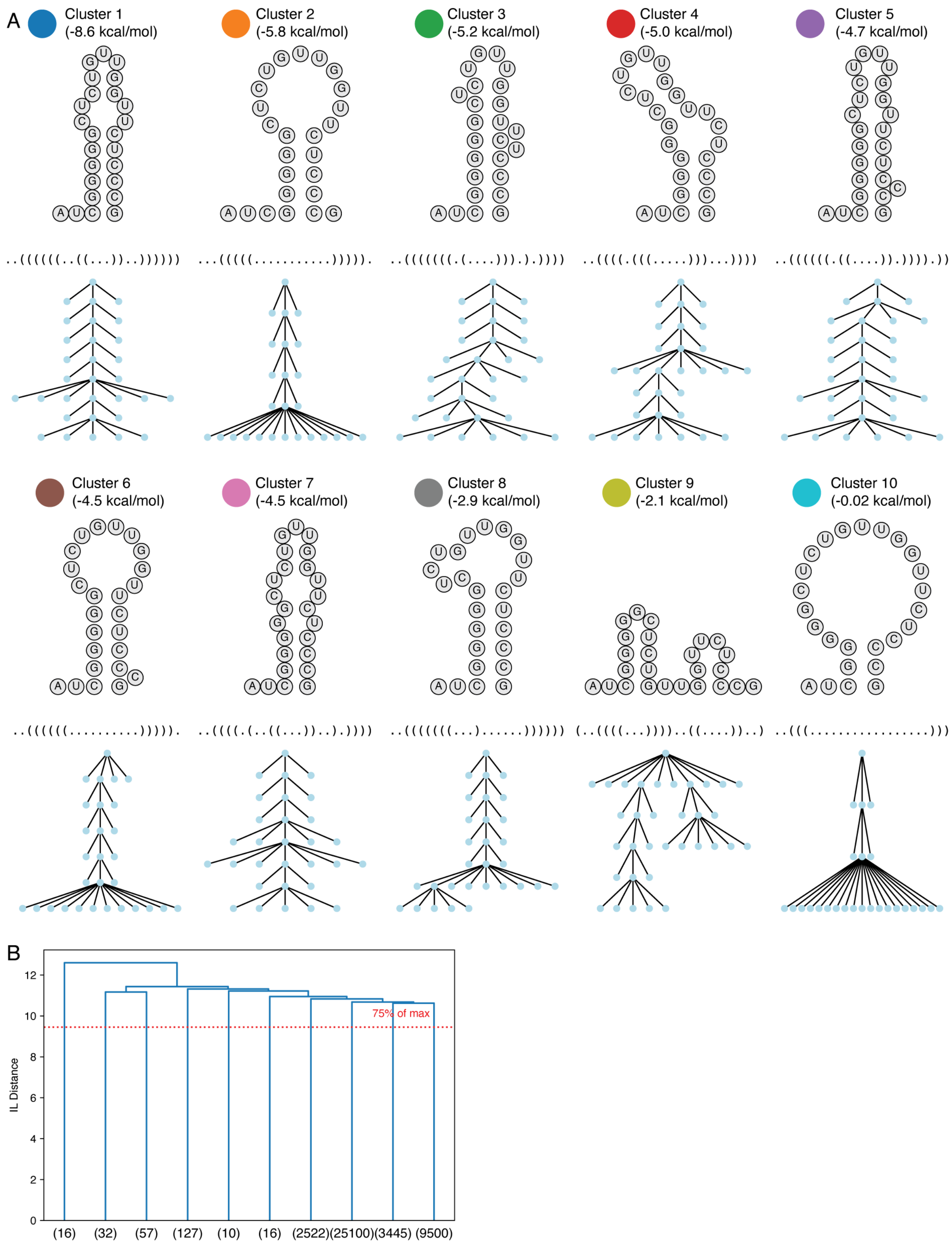

Figure B: **Internal leafset-based structural representation and hierarchical clustering of centroid structures within clusters.** (a): Top ten centroid structures shown in Figure 3, ordered by increasing free energy, alongside their corresponding tree visualisations constructed using the internal leafset (IL) representation. (b): Hierarchical clustering of the structural ensemble using the IL distance metric, with clusters defined by a cutoff at 75% of the maximum pairwise distance.

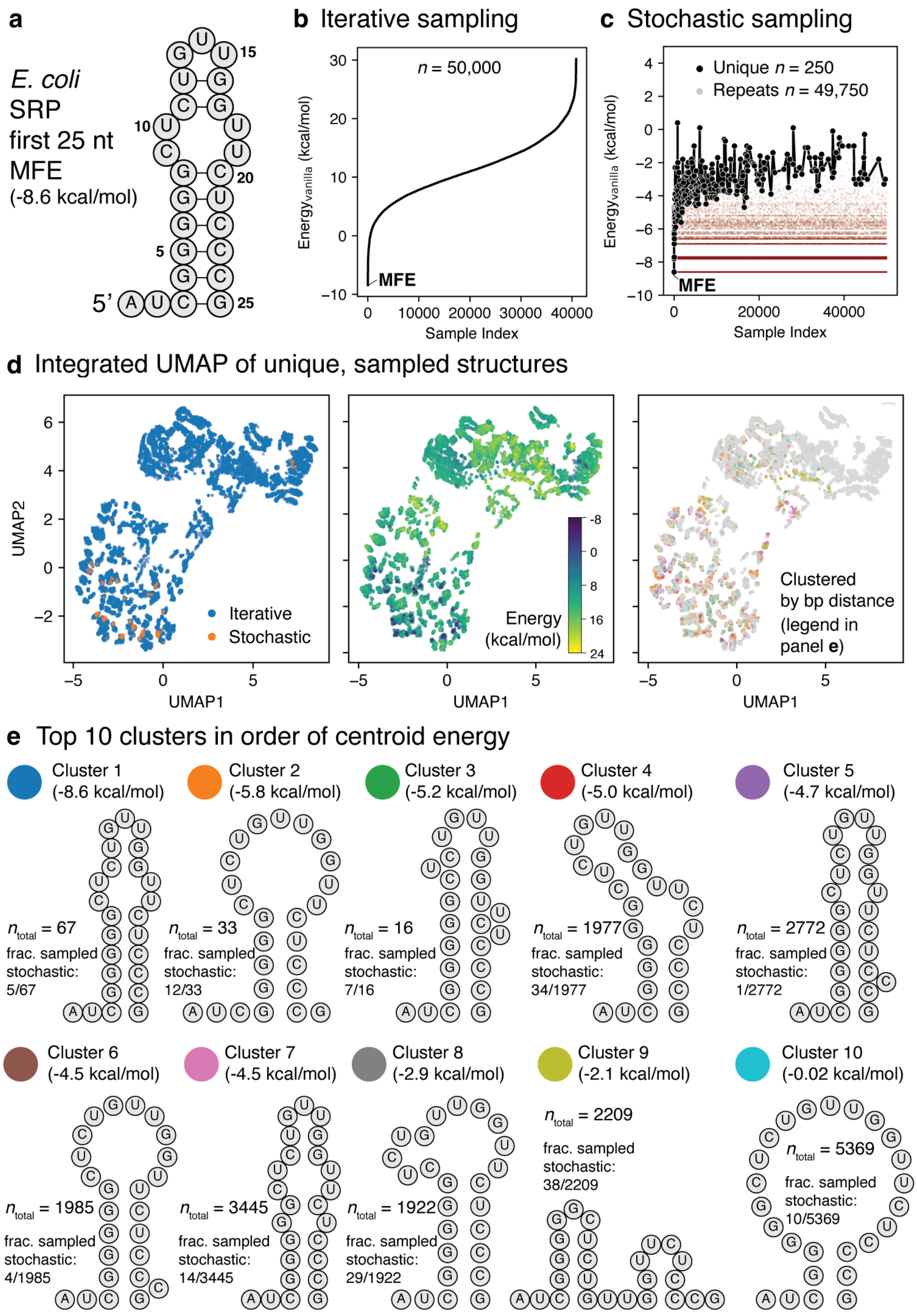

Figure C: **Iterative sampling expands coverage of diverse suboptimal structures.** (a): Predicted minimum free energy (MFE) fold of the first 25 nucleotides of *E. coli* SRP RNA showing a simple hairpin structure. (b): Iterative sampling of this sequence follows a strict order of increasing free energy, producing a sigmoidal profile. (c): Stochastic sampling of the same sequence shows structures generated in a random order, repeatedly sampling low-energy states. (d): Integrated UMAP embedding of all unique sampled structures for the 25-nt SRP RNA (combined superset from iterative and stochastic sampling). Iterative sampling fully encompasses the structures recovered by stochastic sampling. (Left) Points coloured by sampling method (iterative vs. stochastic). (Middle) Points coloured by free energy. (Right) Structures clustered by hierarchical clustering using base-pair distance (Methods); clusters are colour-coded as defined in panel (e). (e): Top 10 clusters ranked by lowest free energy, showing centroid structures, total cluster sizes ($n_{\text{total}}$), and the fraction of each cluster recovered by stochastic sampling.

### B Supplementary Tables

| | RNAstructure | ViennaRNA | memerna-iterative | memerna-persistent |
|---|---|---|---|---|
| RNAstructure | — | — | 0.42 | 0.46 |
| ViennaRNA | — | — | 0.04 | 0.21 |
| memerna-iterative | 2.37 | 23.03 | — | 1.00 |
| memerna-persistent | 2.15 | 4.72 | 1.00 | — |

Table A: Pooled structure count ratio. Values greater than 1 indicate that the row program produced more structures. For ViennaRNA we only included the d2 mode. The sets of structures for each pairwise comparison were slightly different because program execution is limited to a maximum time and memory, which is why the iterative and persistent variants of memerna have different ratios. For the same delta, the iterative and persistent variants of memerna produce the same set of structures.

Table B: Program configuration descriptions

| Identifier | Program | Notes |
|---|---|---|
| RNAstructure | `RNAstructure` 6.5 Fold | Same model as `memerna`, but no sparsification |
| ViennaRNA-d2 | `ViennaRNA` 2.7.2 `RNAsubopt` | d2 option (approximate coaxial stacking) |
| ViennaRNA-all | `ViennaRNA` 2.7.2 `RNAsubopt` | d3 option (full Turner 2004 model) |
| memerna-d2-iterative | `memerna` 0.2 subopt | same as `ViennaRNA` d2, iterative deepening |
| memerna-d2-iterative-lowmem | `memerna` 0.2 subopt | same as `ViennaRNA` d2, iterative deepening, LRU cache |
| memerna-d2-persistent | `memerna` 0.2 subopt | same as `ViennaRNA` d2, persistent |
| memerna-d2-persistent-lowmem | `memerna` 0.2 subopt | same as `ViennaRNA` d2, persistent, LRU cache |
| memerna-all-iterative | `memerna` 0.2 subopt | full Turner 2004 model, iterative deepening |
| memerna-all-iterative-lowmem | `memerna` 0.2 subopt | full Turner 2004 model, iterative deepening, LRU cache |
| memerna-all-persistent | `memerna` 0.2 subopt | full Turner 2004 model, persistent |
| memerna-all-persistent-lowmem | `memerna` 0.2 subopt | full Turner 2004 model, persistent, LRU cache |

| Identifier | Command |
|---|---|
| RNAstructure | DATAPATH=<datapath> AllSub -a <delta> <in> <out> |
| ViennaRNA-d2 | echo <in> \| RNAsubopt --noLP -d2 --sorted -e <delta> > <out> |
| ViennaRNA-all | echo <in> \| RNAsubopt --noLP -d3 --sorted -e <delta> > <out> |
| memerna-all-iterative | subopt --subopt-alg iterative --subopt-sorted --subopt-delta <delta> --backend baseopt --no-ctd-output <in> > <out> |
| memerna-d2-iterative | subopt --subopt-alg iterative --ctd d2 --subopt-sorted --subopt-delta <delta> --backend base --no-ctd-output <in> > <out> |
| memerna-all-iterative-lowmem | subopt --subopt-alg iterative-lowmem --subopt-sorted --subopt-delta <delta> --backend baseopt --no-ctd-output <in> > <out> |
| memerna-d2-iterative-lowmem | subopt --subopt-alg iterative-lowmem --ctd d2 --subopt-sorted --subopt-delta <delta> --backend base --no-ctd-output <in> > <out> |
| memerna-all-persistent | subopt --subopt-alg persistent --subopt-sorted --subopt-delta <delta> --backend baseopt --no-ctd-output <in> > <out> |
| memerna-d2-persistent | subopt --subopt-alg persistent --ctd d2 --subopt-sorted --subopt-delta <delta> --backend base --no-ctd-output <in> > <out> |
| memerna-all-persistent-lowmem | subopt --subopt-alg persistent-lowmem --subopt-sorted --subopt-delta <delta> --backend baseopt --no-ctd-output <in> > <out> |
| memerna-d2-persistent-lowmem | subopt --subopt-alg persistent-lowmem --ctd d2 --subopt-sorted --subopt-delta <delta> --backend base --no-ctd-output <in> > <out> |
| memerna-all-iterative | subopt --subopt-alg iterative --subopt-sorted --subopt-strucs <num> --backend baseopt --no-ctd-output <in> > <out> |
| memerna-d2-iterative | subopt --subopt-alg iterative --ctd d2 --subopt-sorted --subopt-strucs <num> --backend base --no-ctd-output <in> > <out> |
| memerna-all-iterative-lowmem | subopt --subopt-alg iterative-lowmem --subopt-sorted --subopt-strucs <num> --backend baseopt --no-ctd-output <in> > <out> |
| memerna-d2-iterative-lowmem | subopt --subopt-alg iterative-lowmem --ctd d2 --subopt-sorted --subopt-strucs <num> --backend base --no-ctd-output <in> > <out> |
| memerna-all-persistent | subopt --subopt-alg persistent --subopt-sorted --subopt-strucs <num> --backend baseopt --no-ctd-output <in> > <out> |
| memerna-d2-persistent | subopt --subopt-alg persistent --ctd d2 --subopt-sorted --subopt-strucs <num> --backend base --no-ctd-output <in> > <out> |
| memerna-all-persistent-lowmem | subopt --subopt-alg persistent-lowmem --subopt-sorted --subopt-strucs <num> --backend baseopt --no-ctd-output <in> > <out> |
| memerna-d2-persistent-lowmem | subopt --subopt-alg persistent-lowmem --ctd d2 --subopt-sorted --subopt-strucs <num> --backend base --no-ctd-output <in> > <out> |

Table C: Suboptimal folding run commands

| Description | Command |
|---|---|
| memerna compilation command | poetry run python -m rnapy.run build -kind release -energy-precision 1 |
| random dataset creation | poetry run python -m rnapy.run generate-random-dataset -count-per-size 10 -dataset-name random_subopt 20 30 50 80 100 150 200 250 300 400 500 800 1000 1250 1500 1750 2000 2500 3000 |
| Benchmark invocation | poetry run python -m rnapy.run run-subopt-perf -output-dir /bin/subopt_out -dataset random_subopt -num-tries 1 -time-limit-seconds 1800 -memory-limit-bytes 8589934592 -cpu-affinity 1 |

Table D: Benchmark invocation commands

| Key | Value |
| --- | --- |
| RNA lengths, main benchmark (Figure 5, Table 5, Table 6, Supplementary Figure A, Supplementary Table A) | 100, 1000 |
| RNA lengths, scaling statistics (Figure 4) | 20, 30, 50, 80, 100, 150, 200, 250, 300, 400, 500, 800, 1000, 1250, 1500, 1750, 2000, 2500, 3000 |
| Deltas ($\delta$) | 0, 0.1, 0.2, 0.3, 0.4, 0.5, 0.6, 0.7, 0.8, 0.9, 1, 1.2, 1.4, 1.5, 1.8, 2, 3, 4, 5, 6, 7, 8, 9, 10, 15, 20, 25, 30, 35, 45, 60, 75, 100 |
| Number of structures | 1, 10, 100, 1000, 10000, 100000, 1000000, 10000000, 100000000, 1000000000 |

Table E: Benchmark data parameters

### C Supplementary Sections

#### C.1 Existing algorithms

##### C.1.1 Zuker's suboptimal folding method

Zuker's suboptimal folding method [1] is based on finding the MFE traceback multiple times, once with each possible base pair forced to be part of the structure. We describe the algorithm in Algorithm A.

Briefly, it selects base pairs one at a time and computes the best structure that includes that base pair. The best structure that includes a base pair $(i, j)$ is found by computing the traceback of $V(i, j)$ and $V(j, i)$, where $V(i, j)$ is the energy of the interior MFE structure spanning from $i$ to $j$ with a base pair at $(i, j)$, and $V(j, i)$ is the energy of the exterior MFE structure outside of the span from $i$ to $j$, inclusive. The complexity is described in Supplementary Table F.

This algorithm is heuristic in the sense that it cannot exhaustively output all suboptimal structures.

| **Metric** | **Complexity** |
|---|---|
| Time | $O(N^3)$ for the DP, $O(N^4)$ for the tracebacks |
| Time per suboptimal structure | $O(N^2)$ |
| Time per suboptimal structure (amortised) | $O(N^2)$ |
| Memory | $O(N^2)$ for the DP, $O(N)$ for the tracebacks |
| Memory per suboptimal structure | $O(N)$ |
| Memory per suboptimal structure (amortised) | $O(1)$ |

Table F: Time and memory complexity of Zuker's suboptimal folding method.

**Algorithm A** Pseudocode describing Zuker's suboptimal folding method

```
function SuboptZuker(r, δ)
    mfe_dp_tables ← MfeFold(r)
    for all possible base pairs (i, j) do
        if V(i, j) + V(j, i) is within δ then
            Save Traceback(mfe_dp_tables) given (i, j) is paired
        end if
    end for
end function
```

##### C.1.2 Algorithm of Wuchty *et al.*

The algorithm of Wuchty *et al.* applies the Waterman and Byers scheme [2] to RNA folding. They define a partial structure as a stack of sequence substrings and a set of base pairs (not including CTDs, as this algorithm predated those). A complete structure is a partial structure whose stack is empty. They define a refinement as a partial structure that came from another one (they have a more formal definition of this).

**Algorithm B** Pseudocode describing the algorithm of Wuchty *et al.*

```
struct PartialStructure
    unexp: stack of unexpanded states, e.g. {(7, 10, P), (2, 4, U)}
    base_pairs: set of base pairs, e.g. {(1, 5), (6, 11)}
    delta: energy difference from MFE
end struct
function SuboptWuchty(r, δ)
    N ← length of r
    mfe_dp_tables ← MfeFold(r)
    stack ← [PartialStructure{unexp ← (1, N, EXT), base_pairs ← ∅, delta ← 0}]
```

| Metric | Complexity |
|---|---|
| Time | $O(N^3)$ for the DP, $O(SN^3)$ for the tracebacks |
| Time per suboptimal structure | $O(N^3)$ |
| Memory | $O(N^2)$ for the DP, $O(SN^2)$ for the tracebacks |
| Memory per suboptimal structure | $O(N^2)$ |

Table G: Upper bound on time and memory complexity of the algorithm of Wuchty *et al.*

```
    while stack is not empty do
        partial ← pop from stack
        if partial.unexp is empty then                    ▷ Partial structure is complete
            save partial
            continue
        end if
        next_unexp ← pop from partial.unexp
        for all expansion in EXPAND(mfe_dp_tables, next_unexp) do
            if partial.delta + expansion.delta ≤ δ then
                new_partial ← partial with expansion applied
                push new_partial onto stack
            end if
        end for
    end while
end function
```

The algorithm itself does a depth-first search (DFS) on the expansion tree, which stores the entire partial structure in each node. At each node it considers every partial structure it can refine from the current node's partial structure. The refined partial structures whose energy is not more than $\delta$ from the MFE are all pushed onto the stack. When the DFS reaches a fully completed suboptimal structure, it saves it.

The algorithm of Wuchty *et al.* does not provide suboptimal structures in sorted order, and does not support generating structure-by-structure — all structures within $\delta$ are generated in one go. It also must enumerate every expansion at each node in the expansion tree, since they are not sorted so there can be no early stopping.

The complexity analysis for Wuchty-like suboptimal folding algorithms is non-trivial — there is no time or memory complexity given in the original paper [3]. Since there is shared work at internal nodes for each structure, it is difficult to analyse on a per structure basis. It is also tricky to write down the complexity in terms of $\delta$ as it requires knowing how many suboptimal structures will be produced. The analysis in terms of the total number of structures produced, $S$, is also difficult for Wuchty-like algorithms if there is work done for structures not enumerated. For example, at each node the algorithm of Wuchty *et al.* enumerates and performs an $O(N)$ copy for each expansion even if that expansion will not be used to generate a structure.

We place an upper bound on the time and memory usage in Supplementary Table G. The worst case time for one structure is $O(N^3)$, since by Corollary C.1.2 there are $O(N)$ nodes along the path generating it, and at each one looks at $O(N)$ expansions (Corollary C.1.3) and does $O(N)$ copying work for each (Corollary C.1.1). It follows that the worst case total time for the DFS is $O(SN^3)$, although this is not a tight bound as much of the work is shared between structures. By a similar argument the memory is bounded by $O(N^2)$ per suboptimal structure and $O(SN^2)$ total.

##### C.1.3 Stochastic sampling

Stochastic sampling is a method to sample a random structure weighted by its probability in the Boltzmann thermodynamic ensemble of structures. This can be done using the algorithm of Ding and Lawrence [4], with Ponty's improvement [5] improving the worst case runtime from $O(N^2)$ to $O(N \log N)$ and the average case runtime from $O(N\sqrt{N})$ to $O(N \log N)$. We have described it in Algorithm C.

Ponty's improvement is to the selection. For example, it iterates through the splits for multi-loop decomposition in a special order: alternating left and right, going inwards. When the selection algorithm needs to take a lot of steps to select the next pivot, the two child states become a more even split of the parent state. This avoids the pathological case of taking $N$ time to make a split where the left child state's size is still close to size $N$. Repeating this pathological step leads to $O(N^2)$ total time spent in selection. Ponty's improvement brings this down to $O(N \log N)$ by "allocating" even splits to when split selection is slow and uneven splits to when split selection is fast.

**Algorithm C** Pseudocode describing stochastic sampling

```
struct PartialStructure
    unexp: stack of unexpanded states, e.g. {(7, 10, P), (2, 4, U)}
    base_pairs: set of base pairs, e.g. {(1, 5), (6, 11)}
end struct
function StochasticSampling(r)
    N ← length of r
    partition_dp_tables ← Partition(r)
    partial ← {unexp ← (1, N, EXT), base_pairs ← ∅}        ▷ A single PartialStructure
    while partial.unexp is not empty do
        next_unexp ← pop from partial.unexp
        expansion ← select from Expand(partition_dp_tables, next_unexp) weighted by their
probability using partition_dp_tables
        apply expansion to partial in place
    end while
    return partial
end function
```

#### C.2 Algorithmic details

##### C.2.1 Complexity bounds on traceback and partial structures

**Lemma C.1.** *Traceback time is worst-case $O(N^2)$.*

*Proof.* The traceback processes $O(N)$ states. The states are limited to $O(N)$ because each state $S$ covers some area $(st, en)$ which maps to two states replacing $S$ with areas $(st', en')$ and $(st'', en'')$ where both of these are fully contained within the span $(st, en)$, non-overlapping, and non-empty. This process can only happen $O(N)$ times until every state is reduced to size one. This also means that the size of the traceback stack can never exceed $O(N)$ since the states can only be split $N$ times before everything is of area 1.

At each state which covers an area of size $M$ there are $O(M)$ split points to consider. The worst case is if at each state, it is split into a state with an area of 1 and a state with an area of $M - 1$. This gives $O(N^2)$ time for the traceback in total. □

The stack used for the optimal traceback is roughly equivalent to a single partial structure which is evolved in place. As a consequence of this and Theorem C.1 we have Corollary C.1.1, Corollary C.1.2, and Corollary C.1.3.

**Corollary C.1.1.** *The number of unexpanded states in a partial structure is bounded by $O(N)$.*

**Corollary C.1.2.** *The maximum number of unexpanded states seen while evolving a partial structure to a complete structure is $O(N)$.*

**Corollary C.1.3.** *The maximum number of expansions at a state is $O(N)$.*

##### C.2.2 Structure recovery from persistent data structures

**Algorithm D** Pseudocode describing how to recover a structure from the persistent data structure.

**function** GENERATESTRUCTURE
  **args**
    *delta* ▷ The energy delta from MFE for this structure.
    *idx* ▷ The index of the node in the persistent tree.
  **yields** (*base_pairs*, *energy*) ▷ A generated suboptimal structure and its energy.
  $exp\_idx \leftarrow nodes[idx].parent\_exp\_idx$
  $idx \leftarrow nodes[idx].parent\_idx$
  $base\_pairs \leftarrow \emptyset$
  **while** $idx \neq -1$ **do**
    $to\_exp \leftarrow nodes[idx].to\_exp$
    $exp \leftarrow$ INCREMENTALEXPAND$(mfe_dp_tables, to_exp)[exp_idx]$ ▷ Expansion that generated the previous node
    add *exp*'s base pairs to *base_pairs* ▷ If handling CTDs, do that here
    $exp\_idx \leftarrow nodes[idx].parent\_exp\_idx$
    $idx \leftarrow nodes[idx].parent\_idx$
  **end while**
  YIELD$(base_pairs, delta + MFE)$
**end function**

**Algorithm E** Pseudocode describing suboptimal folding using iterative deepening.

**function** SUBOPTITERATIVE
  **args**
    *r* ▷ The RNA primary sequence
    *max_delta* ▷ The maximum energy delta of structures to yield.
    *max_structures* ▷ The maximum number of structures to yield.
    *max_time* ▷ The time budget.
  **yields** (*base_pairs*, *energy*) ▷ A generated suboptimal structure and its energy.
  $mfe\_dp\_tables \leftarrow$ MFEFOLD$(r)$
  **if** outputting up to a given energy delta *max_delta* **then**
    SUBOPTITERATIVESTEP$(r, max_delta, false, \infty)$
  **else if** outputting structures sorted by energy up to *max_delta* **then**
    $delta \leftarrow 0$
    **while** $delta \leq max\_delta$ **do**
      $\_, delta \leftarrow$ SUBOPTITERATIVESTEP$(r, delta, true, \infty)$
    **end while**
  **else if** outputting up to a maximum number of structures *max_structures* **then**
    $delta \leftarrow 0$
    $count \leftarrow 0$
    **while** $count \leq max\_structures$ **do**
      $added\_count, delta \leftarrow$ SUBOPTITERATIVESTEP$(r, delta, true, max_structures - count)$
      $count \leftarrow count + added\_count$
    **end while**
  **else if** outputting until a given time elapses **then**
    $delta \leftarrow 0$
    **while** there is time left **do**
      $\_, delta \leftarrow$ SUBOPTITERATIVESTEP$(r, delta, true, \infty)$
    **end while**
  **end if**
**end function**

**Algorithm F** Pseudocode describing suboptimal folding using persistent data structures.

**function** SUBOPTPERSISTENT
  **args**
    *r* ▷ The RNA primary sequence

*max_delta* ▷ The maximum energy delta of structures to yield.
*max_structures* ▷ The maximum number of structures to yield.
*max_time* ▷ The time budget.
**yields** (*base_pairs*, *energy*) ▷ A generated suboptimal structure and its energy.

```
mfe_dp_tables ← MfeFold(r)
start_state ← the initial state
push Node{exp_idx ← 0, to_exp ← start_state}
push (0, 0) onto pq
if outputting structures up to max_delta then
    while pq is not empty do
        (delta, idx) ← SuboptPersistentStep(r)
        if idx = −1 ∨ delta > max_delta then
            break
        end if
        GenerateStructure(delta, idx)
    end while
else if outputting up to a maximum number of structures max_structures then
    count ← 0
    while pq is not empty ∧count < max_structures do
        (delta, idx) ← SuboptPersistentStep(r)
        if idx = −1 then
            break
        end if
        count ← count + 1
        GenerateStructure(delta, idx)
    end while
else if outputting until a given time elapses then
    while pq is not empty ∧ there is time left do
        (delta, idx) ← SuboptPersistentStep(r)
        if idx = −1 then
            break
        end if
        GenerateStructure(delta, idx)
    end while
end if
end function
```

##### C.2.3 Complexity of iterative-strucs

**Time.** It takes $O(N^3)$ for the DP and to generate the expansions, $O(K) = O(S)$ for the sampling, or $O(KN) = O(SN)$ if copying each structure.

For a particular iterative deepening step, we only expand internal nodes which lead to at least one terminal node by pruning according to delta. The path down the expansion tree is at most $O(N)$ (Corollary C.1.2). This means that the number of internal nodes is limited by $O(SN)$, although it appears $O(S)$ per our empirical results. For the exponential regime, we can ignore multiple iterative deepening steps, since the cost will be dominated by the last step. Assuming that $O(S)$ is the number of internal nodes, we produce a structure (which takes $O(N)$ copying time (Corollary C.1.1)) every $O(1)$ internal nodes.

**Memory.** It takes $O(N^3)$ to store the expansions (lowered to $O(N^2)$ if using LRU cache), $O(N^2)$ for the DP table, $O(1)$ memory for global state, and $O(N)$ memory for the DFS stack (Corollary C.1.2). Notably, the memory usage does not depend on the number of structures produced.

##### C.2.4 Complexity of iterative-delta

Per our empirical results while in the exponential regime, the number of structures $S = O(k^\delta)$. So we just replace $S$ with $k^\delta$ to get the complexities for delta suboptimal folding.

##### C.2.5 Complexity of persistent-strucs

**Time.** It takes $O(N^3)$ for the DP and to generate the expansions, $O(K \log K) = O(S \log S)$ for the sampling, or $O(NK + K \log K) = O(SN + S \log S)$ if reconstructing each structure.

It takes $O(\log K) = O(\log S)$ time to select the next node to process. Since we estimate $K = O(S)$ by a similar argument it takes $O(S \log S)$ time to find all structures, and $O(SN)$ time to reconstruct them all from the persistent data structure (Corollary C.1.2).

**Memory.** Since this variant performs a PFS rather than a DFS we need to store each node of the tree, which takes an extra $K = O(S)$ memory.

##### C.2.6 Complexity of persistent-delta

Similarly replace and simplify the expression using $S = O(k^\delta)$.

#### C.3 Iterative sampling increases diversity and massively expands coverage of the energy landscape

We evaluated the behaviour of iterative sampling implemented in `memerna` using a 25-nt sequence at the 5′ end of the *E. coli* SRP RNA (Supplementary Figure Ca) [6], [7].

**Ordered enumeration of structures** Consistent with its design, `memerna` enumerates structures in strict order of increasing free energy, producing a monotonic trajectory through the landscape (Supplementary Figure Cb). The resulting energy profile exhibits an inverse sigmoidal shape: an initial rapid rise from the MFE, followed by a plateau of near-degenerate conformations, and then progression into higher-energy states. In contrast, stochastic Boltzmann sampling generates structures in effectively random order and repeatedly resamples low-energy conformations (Supplementary Figure Cc). After 50,000 draws, only ∼250 unique structures are recovered, indicating substantial redundancy. Moreover, stochastic sampling remains confined to moderately stable states (<0 kcal/mol), whereas iterative sampling traverses the full accessible energy range. These results confirm that iterative sampling deterministically explores the landscape without redundancy, while stochastic sampling provides limited and biased coverage.

**Clustering reveals expanded coverage from iterative sampling** To assess structural diversity, we analysed the union of structures generated by both methods. After removing duplicates, we computed pairwise distances using the internal leafset (IL) metric [8] and visualised the ensemble using UMAP (Supplementary Figure Cd). Hierarchical clustering identified >20 distinct structural clusters spanning the landscape (Supplementary Figure B). While stochastic sampling recovers representatives from the lowest-energy clusters, it captures only a small fraction of structures within each cluster, indicating limited intra-cluster diversity. In contrast, iterative sampling spans all clusters and systematically enumerates their members.

Cluster-level analysis further highlights this disparity. The lowest-energy cluster is small ($n = 67$), reflecting limited variation around the stable hairpin. Higher-energy clusters are substantially larger (e.g., Cluster 4: $n = 1{,}977$), representing diverse conformational classes. Stochastic sampling captures only a sparse subset of these clusters, often sampling them fewer than ten times, whereas iterative sampling comprehensively covers both low- and high-energy regions of the landscape.

Together, these results demonstrate that iterative sampling dramatically increases structural diversity and enables systematic exploration of conformational space, including higher-energy states that are largely inaccessible under Boltzmann-weighted sampling.

### D Supplementary Methods

#### D.1 Additional Benchmark Details

We ran `ViennaRNA` with the `-noLP` option to match `memerna` and `RNAstructure`. While `memerna` supports outputting structures in non-sorted order and this can be faster, we restricted it to output structures in sorted order to match the other programs. We also built `memerna` using 0.1 kcal/mol precision to match `RNAstructure`.

## D.2 R2D2 v2

R2D2 computes a difference score ($D$, also referred to as `rho_dist`) between a reactivity ($\rho$) vector and a binary structural pairing vector. Details of this metric are described in [9]. As in the original R2D2 work [9], reactivities were capped at 1.0. Paired regions were assigned a weight of 0.8 in the distance calculation, while unpaired regions were weighted 0.2.

For the results shown in Figure 6c–d, version 6.4 of `RNAstructure` (using the `stochastic-smp` and `partition-smp` tools) was used and compared against `memerna` version 0.2.0. Runtime measurements in Figure 2C were obtained on a 2.4 GHz Intel Xeon Silver processor with 16 cores and 64 GB RAM, running as the sole user process on an Ubuntu 18 virtual machine. Reported runtimes include both sampling and constant overhead from parsing reactivity files and other R2D2 preprocessing steps.

Example JSON configuration files used to generate datapoints at $n = 64{,}000$ for Figures 2C and 2D are shown below.

**Distance function** is quantified by comparing the experimental reactivity vector $\rho$, which reports nucleotide accessibility ($\rho_i \geq 0$), to a structural accessibility representation defined by the set of unpaired nucleotides $U$. For each nucleotide position $i \in 1, \ldots, n$, accessibility is encoded using an indicator function $I_U(i)$, where $I_U(i) = 1$ if $i \in U$ (unpaired) and $I_U(i) = 0$ otherwise. For each structure, the distance is computed as weighted sum of absolute differences:

$$D_\alpha(U, \rho) = \alpha \sum_{i \in \bar{U}} |I_U(i) - \rho_i| + (1 - \alpha) \sum_{i \in U} |I_U(i) - \rho_i|,$$

where $\alpha$ adjusts the relative contribution of positions predicted to be paired in the sampled structure. Structures that minimise $D_\alpha$ are selected as best matching the experimental chemical probing accessibility profile.

**Example configurations (Figure 6c–d, $n = 64{,}000$)**

```
{
    "run_name": "111nt_memerna_eq_vanilla",
    "reactivity_file": "~/SRP_data/SRP_EQ/SRP_IDT2_mod_111nt_130_reactivities.tx
    "endcut": 18,
    "sampling_config": {
        "sample_size": 64000,
        "sample_gen": "Memerna",
        "bias": "vanilla"
    }
}

{
    "run_name": "111nt_rnastructure_eq_vanilla",
    "reactivity_file": "~/SRP_data/SRP_EQ/SRP_IDT2_mod_111nt_130_reactivities.tx
    "endcut": 18,
    "sampling_config": {
        "sample_size": 64000,
        "sample_gen": "RNAstructure",
        "bias": "pooled"
```

```
    },
    "free_energy_config": {
        "free_energy_fn": "RNAstructure"
    }
}
```

**Configuration for Supplementary Figure Bc**

```
{
    "run_name": "25nt_rnastructure_eq_vanilla_dups",
    "reactivity_file": "~/SRP_data/SRP_EQ/SRP_IDT2_mod_25nt_44_reactivities.txt"
    "endcut": 18,
    "save_ensembles": true,
    "sampling_config": {
        "sample_size": 50000,
        "allow_duplicates": true,
        "sample_gen": "RNAstructure",
        "bias": "vanilla"
    },
    "free_energy_config": {
        "free_energy_fn": "RNAstructure",
        "report_mfe": true
    }
}
```

**SHAPE-constrained configurations (Figure 7c)**

```
{
    "run_name": "25nt_memerna_eq_shape",
    "reactivity_file": "~/SRP_data/SRP_EQ/SRP_IDT2_mod_25nt_44_reactivities.txt"
    "endcut": 18,
    "save_ensembles": true,
    "sampling_config": {
        "sample_size": 150000,
        "sample_gen": "Memerna",
        "bias": "shape",
        "shape_slope": 3.5,
        "shape_intercept": -0.9
    },
    "free_energy_config": {
        "report_mfe": true
    }
}
```

```
{
    "run_name": "25nt_rnastructure_eq_pooled",
    "reactivity_file": "~/SRP_data/SRP_EQ/SRP_IDT2_mod_25nt_44_reactivities.txt"
    "endcut": 18,
    "save_ensembles": true,
    "sampling_config": {
        "sample_size": 150000,
        "sample_gen": "RNAstructure",
        "bias": "pooled",
        "shape_slope": 3.5,
        "shape_intercept": -0.9
    },
```

```
    "free_energy_config": {
        "free_energy_fn": "RNAstructure",
        "report_mfe": true
    }
}
```

SHAPE slope and intercept parameters were used to convert SHAPE reactivities ($\rho$ values) into pseudo-free energy biases ($pf$) for `memerna` according to

$$pf = m \cdot \ln(\rho + 1) + b.$$

The parameters $m = 3.5$ and $b = -0.9$ were empirically determined, inspired by prior work [10].

##### D.3 Pairwise structural distance using Internal-Leafset (IL) distance

We computed pairwise distances between sampled RNA secondary structures using the Internal-Leafset (IL) distance [8], a tree-based metric that compares internal loop topology and nesting structure. This was compared to structural ensembles (e.g., with versus without chemical probing). Unlike base-pair distance, IL distance is sensitive to loop configuration and hierarchical structural organisation.

We represented secondary structures in dot–bracket notation. For each transcript length, we enumerated all unique structures identified by iterative sampling and computed the full pairwise IL distance matrix using the `rnadist` implementation [8]. We parallelised distance calculations for scalability.

For an ensemble of N structures, this produced a symmetric $N \times N$ distance matrix used for downstream clustering and manifold embedding analyses.

##### D.4 Hierarchical clustering of sampled structures

We performed agglomerative hierarchical clustering using the precomputed IL distance matrix. We computed linkage relationships using `SciPy`'s hierarchical clustering implementation and stored them as a linkage matrix $Z$, where each row encodes a merge event and its associated inter-cluster distance.

We obtained cluster assignments by cutting the dendrogram at a distance threshold corresponding to 75% of the maximum linkage height:

$$t = 0.75 \times \max(Z[:, 2]).$$

We extracted flat clusters using the distance criterion:

$$\texttt{fcluster}(Z, t, \text{criterion="distance"}).$$

This procedure partitions the ensemble into clusters whose maximum internal linkage distance does not exceed the specified threshold. We assigned cluster labels to each structure and used them to quantify cluster occupancy, dominant structural basins, and condition-specific enrichment. Because IL distances reflect nucleotide-level pairing differences, hierarchical clustering groups structures into basins of similar pairing topology, providing an interpretable decomposition of the sampled conformational landscape.

##### D.5 UMAP embedding of structural space

To visualise high-dimensional structural relationships, we embedded the $IL$ distance matrix into two dimensions using Uniform Manifold Approximation and Projection (UMAP). We supplied the precomputed pairwise distance matrix $D$ directly to the UMAP reducer with metric set to "precomputed".

We ran UMAP with the following parameters:

- `n_neighbors = 20`, defining the size of the local neighbourhood used to construct the manifold approximation.

- `min_dist = 0.0`, allowing tight packing of similar structures in the embedding space.
- `metric = "precomputed"`, indicating that IL distances were provided explicitly.
- `random_state = 0`, ensuring reproducibility of the embedding.

We obtained the embedding coordinates as

$$Y = \mathrm{UMAP}(D),$$

yielding a two-dimensional representation in which structures with small IL distances are positioned proximally. This embedding preserves local structural similarity while providing a global visualisation of conformational basins and transitions.

We overlaid cluster labels derived from hierarchical clustering onto the UMAP embedding to relate discrete structural basins to continuous manifold geometry. Together, hierarchical clustering and UMAP provide complementary views of ensemble organisation: clustering identifies discrete conformational groups, whereas UMAP reveals the geometric structure and relative separation of these basins within the full structural landscape.

##### D.6 Boltzmann-weighted centroid selection within structural clusters

We defined representative structures for each structural cluster using a Boltzmann-weighted medoid criterion, following the centroid framework introduced by [11]. Rather than selecting the minimum free energy structure, this approach identifies the structure that minimises the expected distance to all other structures in the cluster, thereby representing the thermodynamic ensemble.

For a cluster containing $m$ structures with indices $\{i_1, \ldots, i_m\}$, let $D_{ij}$ denote the precomputed pairwise IL distance between structures $i$ and $j$, and let $E_j$ denote the free energy of structure $j$. We computed Boltzmann weights within each cluster as

$$w_j = \frac{\exp\left[-(E_j - E_{\mathrm{min}})/(RT)\right]}{\sum_{k=1}^{m} \exp\left[-(E_k - E_{\mathrm{min}})/(RT)\right]},$$

where $E_{\mathrm{min}}$ is the minimum energy within the cluster, $R$ is the gas constant (in $\mathrm{kcal\,mol^{-1}\,K^{-1}}$), and $T$ is absolute temperature. Subtracting $E_{\mathrm{min}}$ ensures numerical stability.

For each candidate structure $i$ in the cluster, we computed its Boltzmann-weighted expected distance to all other cluster members:

$$\mathrm{score}(i) = \sum_{j=1}^{m} w_j \, D_{ij}.$$

We defined the centroid (medoid) as the structure minimising this score. This procedure selects the structure with minimal ensemble-averaged structural distance, providing a representative topology that reflects thermodynamic weighting rather than energy alone.

We assigned clusters containing fewer than two structures their sole member as centroid. For each cluster, we additionally computed the effective sample size (ESS) of the Boltzmann weights,

$$\mathrm{ESS} = \frac{1}{\sum_{j=1}^{m} w_j^2}.$$

to quantify weight concentration within the cluster.

This centroid definition yields a thermodynamically weighted representative structure for each conformational basin and is consistent with established centroid concepts in RNA secondary structure analysis [11].

##### D.7 Jensen-Shannon divergence between ensembles

To quantify structural divergence between equilibrium and cotranscriptional folding, we defined an ensemble-level metric based on the Jensen–Shannon divergence (JSD). Unlike distance-based selection metrics applied to individual structures, JSD compares full probability distributions over a shared discrete structure space.

For each transcript length, exhaustive SHAPE-constrained iterative sampling generated a set of unique secondary structures, denoted $\mathcal{S}$. We used the union of structures identified under equilibrium and cotranscriptional conditions to define a common structure space. For each structure $s \in \mathcal{S}$, we computed condition-specific Boltzmann probabilities:

$$P_{\text{equil}} = \{p_{\text{equil}}(s)\}_{s\in\mathcal{S}}, \quad P_{\text{cotrans}} = \{p_{\text{cotrans}}(s)\}_{s\in\mathcal{S}}.$$

We calculated structure probabilities as

$$p(s) = \frac{\exp\left(-\Delta G(s)/kT\right)}{Q} \tag{A}$$

where $\Delta G(s)$ is the reactivity-constrained free energy of structure $s$, $k$ is the Boltzmann constant, $T$ is temperature in Kelvin, and

$$Q = \sum_{s\in\mathcal{S}} \exp\left(-\Delta G(s)/kT\right) \tag{B}$$

is the partition function over the enumerated structure set. We computed free energies using nearest-neighbour thermodynamic parameters augmented with condition-specific pseudo-energy restraints derived from experimental probing data.

We defined the Jensen-Shannon divergence between the two distributions as

$$\text{JSD}(P_{\text{equil}} \parallel P_{\text{cotrans}}) = \frac{1}{2} D_{\text{KL}}(P_{\text{equil}} \parallel M) + \frac{1}{2} D_{\text{KL}}(P_{\text{cotrans}} \parallel M) \tag{C}$$

where

$$M = \frac{1}{2}\left(P_{\text{equil}} + P_{\text{cotrans}}\right)$$

and $D_{\text{KL}}$ denotes the Kullback-Leibler divergence:

$$D_{\text{KL}}(P \parallel Q) = \sum_{s\in\mathcal{S}} P(s) \log_2 \frac{P(s)}{Q(s)} \tag{D}$$

We computed logarithms in base 2, yielding JSD values bounded between 0 and 1. A JSD of 0 indicates identical structural ensembles, whereas a value approaching 1 indicates maximally distinct, non-overlapping distributions.

By operating on thermodynamically weighted distributions defined over the same deterministically enumerated structure space, this framework enables direct and quantitative comparison of equilibrium and cotranscriptional folding landscapes at each transcript length.
